# HNF1A recruits UTX to activate a differentiation program that suppresses pancreatic cancer

**DOI:** 10.1101/690552

**Authors:** Mark Kalisz, Edgar Bernardo, Anthony Beucher, Miguel Angel Maestro, Natalia del Pozo, Irene Millán, Lena Haeberle, Martin Schlensog, Sami Alexander Safi, Wolfram Trudo Knoefel, Vanessa Grau, Matías de Vas, Karl B. Shpargel, Eva Vaquero, Terry Magnuson, Sagrario Ortega, Irene Esposito, Francisco X. Real, Jorge Ferrer

## Abstract

Defects in transcriptional regulators of pancreatic exocrine differentiation have been implicated in pancreatic tumorigenesis, but the molecular mechanisms are poorly understood. The locus encoding the transcription factor HNF1A harbors susceptibility variants for pancreatic ductal adenocarcinoma (PDAC), while *KDM6A*, encoding the histone demethylase UTX, carries somatic mutations in PDAC. Here, we show that pancreas-specific *Hnf1a* null mutations phenocopy *Utx* deficient mutations, and both synergize with *Kras^G12D^* to cause PDAC with sarcomatoid features. We combine genetic, epigenomic and biochemical studies to show that HNF1A recruits UTX to genomic binding sites in pancreatic acinar cells. This remodels the acinar enhancer landscape, activates a differentiation program, and indirectly suppresses oncogenic and epithelial-mesenchymal transition genes. Finally, we identify a subset of non-classical PDAC samples that exhibit the *HNF1A/UTX*-deficient molecular phenotype. These findings provide direct genetic evidence that HNF1A-deficiency promotes PDAC. They also connect the tumor suppressive role of UTX deficiency with a cell-specific molecular mechanism that underlies PDAC subtype definition.

## Introduction

Pancreatic ductal adenocarcinoma (PDAC) is a leading cause of cancer mortality ^1^. The incidence of PDAC is rising, yet current chemotherapies are generally ineffective ^2^. Genomic analysis of PDAC has identified almost universal driver mutations in *KRAS, TP53, SMAD4* and *CDKN2A*, among a long list of loci that show recurrent somatic mutations and structural variations ^3–7^. A small subset of tumors is caused by germ-line mutations in DNA-repair genes ^5, 8^, whereas GWAS have identified dozens of common variants that impact PDAC susceptibility ^9, 10^. Genetic studies have therefore uncovered leads that promise to define molecular targets for future precision therapies.

Up to 18% of PDAC tumors carry mutations in *KDM6A* ^5^, which encodes UTX, a component of the MLL/COMPASS transcriptional co-regulatory complex ^11^. UTX catalyzes demethylation of histone H3K27me3, a modification associated with Polycomb-mediated repression ^12–14^. Most somatic pathogenic *UTX* mutations are likely to result in a loss of function, and mouse genetic studies have shown that *Utx* and *Kras* mutations cooperate to promote PDAC ^15, 16^. How UTX is recruited to its genomic targets in pancreatic cells, and the direct mechanisms through which it controls PDAC-relevant genetic programs, is still poorly understood ^17^.

There is increasing evidence that the transcriptional regulation of differentiated pancreatic exocrine cells is tightly linked to PDAC development and subtype definition ^18–22^. Little is known, however, about the underlying molecular underpinnings. We have examined HNF1A, a homeodomain transcriptional regulator of liver, gut, kidney as well as pancreatic acinar and endocrine cells, which has been proposed to act as a candidate pancreatic tumor suppressor ^22–25^. Human heterozygous *HNF1A* loss of function mutations cause diabetes, in part because *HNF1A* promotes pancreatic β cell proliferation, and mouse *Hnf1a* mutations prevent the formation of large T antigen-driven β cell tumors ^26^. The function of HNF1A, however, is cell-type specific ^26^, and both co-expression network analysis of PDAC samples as well as *in vitro* studies suggest that *HNF1A* has a tumor suppressive function in pancreatic exocrine cells ^23, 24^ Furthermore, GWAS have shown that the *HNF1A* locus contains one of the strongest PDAC genetic association signals ^10, 27^ Despite these observations, there is currently no direct mouse or human genetic evidence to incriminate *HNF1A* deficiency in PDAC.

Here, we combine mouse genetic knock-outs, transcriptomics and genome binding studies to show that HNF1A is a major determinant for the recruitment of UTX to its genomic targets in acinar cells. This recruitment remodels the enhancer landscape of acinar cells and activates a broad epithelial cell transcriptional program, which in turn indirectly suppresses tumor suppressor pathways. We demonstrate that *Hnf1a* inactivation promotes Kras-induced PDAC, and partially phenocopies morphological features of Utx-driven PDAC. Finally, we define a subset of human tumors that exhibit *HNF1A/UTX*-deficient transcriptional programs. These findings, therefore, provide a molecular mechanism that connects the tumor suppressive functions of UTX and pancreatic differentiation transcription factors.

## Results

### Hnf1a-deficiency promotes Kras-induced oncogenesis

To directly test the role of *Hnf1a* in pancreatic carcinogenesis, we created a conditional *Hnf1a* loss of function allele (*Hnf1a^LOxP^*) (**Supplementary Figure 1a**) and used a *Pdx1^Cre^* transgene to delete *Hnf1a* in all pancreatic epithelial lineages (hereafter referred to as *Hnf1a^pKO^* mice, **Supplementary Figure 1b**). HNF1A is normally expressed in pancreatic acinar and endocrine cells, but not in duct cells ^28^, and *Hnf1a*^pKO^ mice showed disrupted HNF1A expression in both acinar and endocrine cells (**Supplementary Figure 1c**). As expected from previous studies of *Hnf1a* germ-line null mutants, this did not produce gross defects in pancreas organogenesis, although acinar cells displayed signs of markedly increased proliferation ^25, 29–31^ (**Figure 1a**).

**Figure 1.**
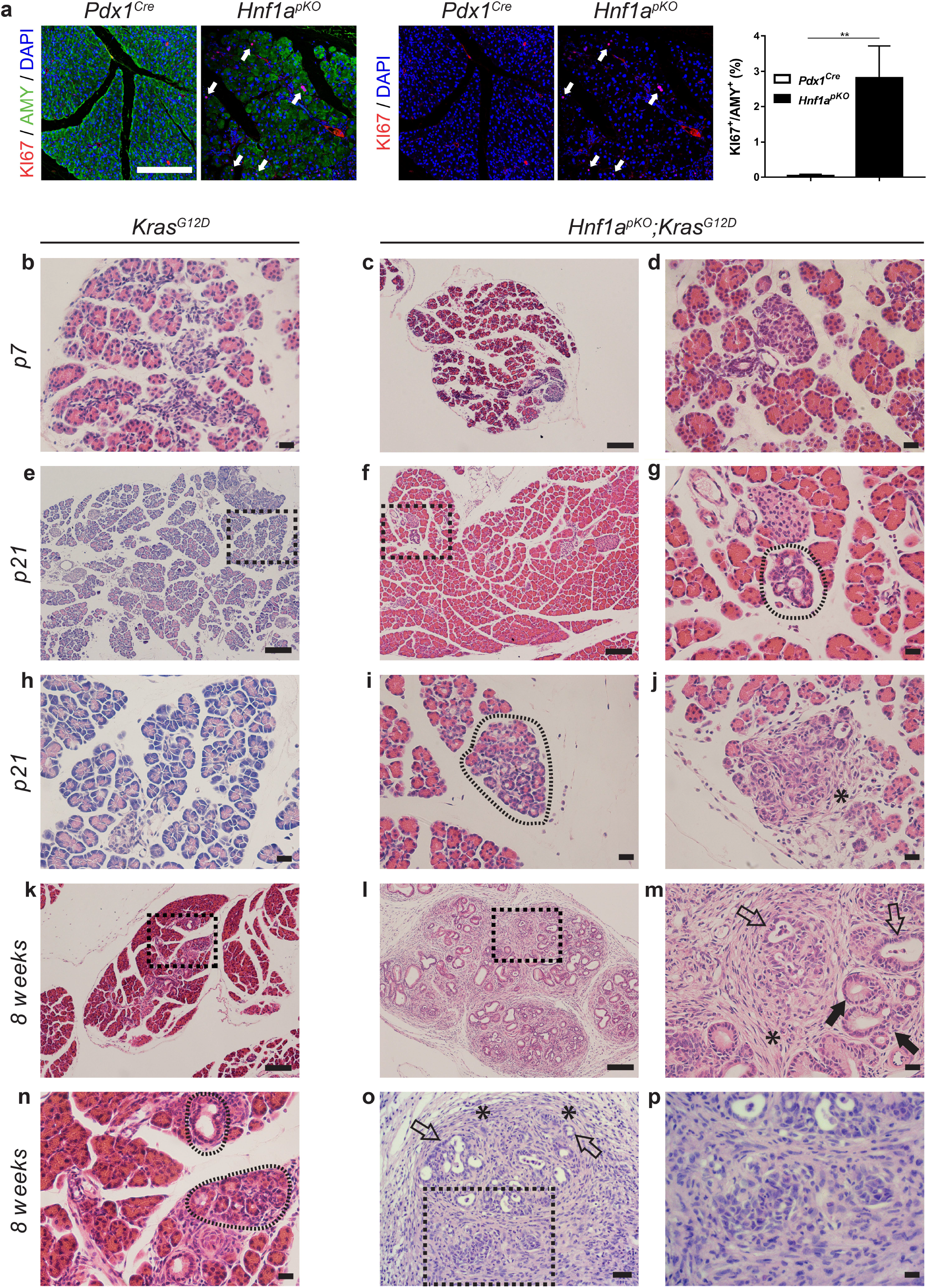
HNF1A-deficiency leads to increased proliferation and promotes Kras-induced oncogenesis. **a**, Representative immunofluorescence images and quantifications showing that 3 months old *Hnf1a^pKO^* mice have increased number of KI67^+^ (red) acinar cell nuclei costaining with DAPI (blue) and Amylase (green). Arrows point to KI67+ acinar cells in *Hnf1a^pKO^* mouse. Scale bar is 200 μm. Acinar proliferation was represented as the average of the Ki67+/Amylase+ cell ratio. Quantifications were performed on 3 random fields from 3 *Pdx1^Cre^* and 3 *Hnf1a^pKO^* mice. Student’s *t* test p<0.01. Representative H&E stainings of pancreata from *Kras^G12D^* and *Hnf1a^pKO^*;*Kras^G12D^* mice. **b-d**, *Kras^G12D^* and *Hnf1a^pKO^*;*Kras^G12D^* mice have normal morphology at 7 days, **e-j**, at 21 days *Hnf1a^pKO^*;*Kras^G12D^* mice show acinar-to-ductal metaplasia (dashed encircled areas) and regions with desmoplastic reaction (asterisk), which are not observed in *Kras^G12D^* mice (**e, h**). **k-p**, At 8 weeks *Kras* pancreas show occasional abnormal ductal structures (dashed encircled areas in n) and **l,m,o,p**, *Hnf1a^pKO^*;*Kras^G12D^* mice present mucinous tubular complexes (black arrows), and more advanced PanINs with luminal budding (open arrows) including foci of spindle cell proliferation (asterisks) and incipient infiltrative growth (black dashed box area on **o**). Black dashed boxes in e,f, k, l and o indicate magnified areas in h, g, n, m and p respectively. Scale bars indicate 100 μm (c,e,f,k,l), 50 μm (o) and 20 μm (b,d,g,h-j,m,n,p).

To determine whether *Hnf1a* interacts with *Kras*-induced carcinogenesis, we created mice with combined conditional *Hnf1a* and *Kras^G12D^* mutations, hereafter referred to as *Hnf1a^pKO^*;*Kras^G12D^* mice (**Supplementary Figure 1d**). In the absence of *Hnf1a* mutant alleles, *Pdx1^Cre^*-induced *Kras^G12D^* activation expectedly gave rise to occasional low-grade PanINs or acinar-to-ductal metaplasia (ADM) lesions by 2 months of age ^32^, (**Figure 1b, e, h, k**, and **n**). *Hnf1a^pKO^*;*Kras^G12D^* mice showed no lesions at 7 days of age (**Figure 1c-d**), yet by weaning they had already developed focal ADM and desmoplastic reactions, which became more prominent as the mice aged (**Figure 1f-g and Figure 1i-j**). Eight-week-old *Hnf1a^pKO^*;*Kras^G12D^* mice additionally showed non-invasive atypical tubular complexes, higher-grade PanINs with luminal budding, desmoplastic reaction and foci of spindle cell (mesenchymal) proliferation, some of which showed incipient infiltrative growth (**Figure 1l-m and Figure1o-p**). These findings indicate that pancreatic *Hnf1a*-deficiency cooperates with *Kras* to promote sarcomatoid forms of PDAC.

### *Hnf1a* activates an acinar differentiation program that inhibits oncogenic programs

To understand how *Hnf1a*-deficiency promotes pancreatic cancer, we examined the transcriptional programs controlled by *Hnf1a* in pancreatic exocrine cells. Genetic lineage tracing studies in mice have shown that, despite the ductal morphology of PDAC, *Kras^G12D^*-induced PDAC can originate from acinar cells that undergo ADM and PanIN, contrasting with intraductal papillary mucinous neoplasms that arise from duct cells ^33, 34^. We therefore used a *Ptf1a^Cre^* allele, which ensured high efficiency recombination of *Hnf1a* in mouse acinar cells, and more limited recombination in endocrine cells (*Ptf1a^Cre^*; *Hnf1a^LoxP/LoxP^*; hereafter referred to as *Hnf1a^aKO^*) (**Supplementary Figure 2a and 2b**). Eight-week-old *Hnf1a^aKO^* mice were normoglycemic, and like *Hnf1a^pKO^* mice showed normal pancreatic histology (**Supplementary Figure 2c**). We profiled transcripts in pancreas from 8-week-old *Hnf1a^aKO^* mice and, despite the normal histology, found profound transcriptional changes (**Figure 2a, Supplementary Table 1**). We observed decreased expression of genes specific to differentiated acinar cells, including *Ptf1a, Pla2g1b, Serpini2* and *Ctrb1* (**Figure 2b and Supplementary Figure 2d**), and increased expression of genes specific to pancreatic mesenchymal cells (**Figure 2b, Supplementary Table 2**). Downregulated genes were enriched in metabolic processes such as inositol phosphate turnover, amino acid metabolism, and protection against oxidative stress, whereas upregulated genes were enriched in annotations associated with the extracellular matrix (collagen formation, ECM-receptor interactions, integrin cell surface interactions) and complement activation (**Figure 2c, Supplementary Table 2**). We also observed activation of known oncogenic programs such as EMT, RAS, PI3K-AKT, STAT3, MAPK, and WNT signaling, cell cycle related pathways, and cholesterol biosynthesis (**Figure 2d, Supplementary Figure 2e, Supplementary Table 2**).

**Figure 2.**
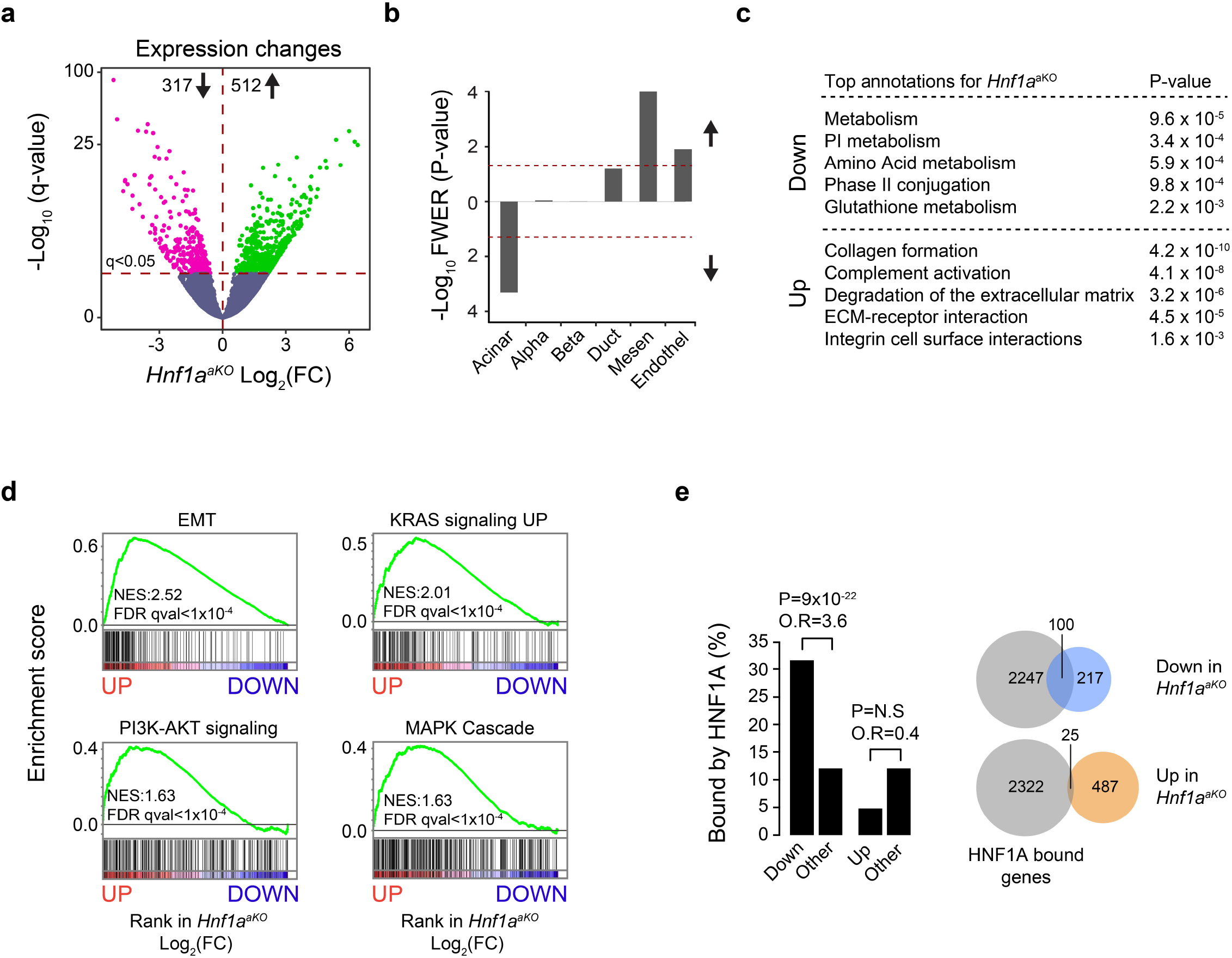
HNF1A regulates gene programs essential to acinar cell identity, metabolism and proliferation. **a**, Fold change (FC) in transcripts in *Hnf1a^aKO^* vs. control pancreas, plotted against significance (-Log_10_ q; genes significant at q<0.05 are shown as colored dots above the horizontal line). **b**, GSEA showing that genes specific to differentiated acinar cells were downregulated in *Hnf1a^aKO^* pancreas, but not genes specific to islets or duct cells. Upregulated genes were enriched in genes specific to mesenchymal cells. Lineage-enriched genes were obtained from Muraro et al. ^1^. **c**, Top functional annotations for differentially expressed genes in *Hnf1a^aKO^* pancreas. **d**, GSEA revealed that *Hnf1a^aKO^* pancreas showed increased transcripts involved in oncogenic pathways such as EMT, MAPK, KRAS, PI3K-AKT. **e**, HNF1A promotes transcriptional activation of direct target genes. Left: HNF1A bound-genes were enriched among genes that showed downregulation in *Hnf1a* mutants, but not among upregulated genes. P-values and odds ratios (O.R.) calculated by Fisher’s exact test. Right: Venn diagrams showing overlap of HNF1A-bound genes with genes that were downregulated and upregulated in *Hnf1a* mutant pancreas.

To assess which of these transcriptional changes reflects a direct function of HNF1A in acinar cells, we profiled genome-wide binding sites of HNF1A in adult pancreas, as well as H3K27 acetylation to mark active enhancers and promoters (**Supplementary Figure 2f**). HNF1A-bound genomic regions had canonical HNF1 recognition sequences in 405 out of the top 500 most significant binding sites (81%), confirming the specificity of the assay (**Supplementary Figure 2g**), and they were expectedly enriched in enhancers and promoters (**Supplementary Figure 2h**). HNF1A binding was specifically enriched amongst genes that showed downregulation in *Hnf1a^aKO^* pancreas (odds ratio = 3.6 relative to all other genes, *P* = 10^−22^, **Figure 2e**). This was consistent with HNF1A acting as an essential positive regulator of some of the genes to which it binds. HNF1A binding was, however, not enriched in genes that were upregulated in *Hnf1a^aKO^* pancreas, compared with active genes that did not show altered regulation (odds ratio = 0.4, P=N.S; **Figure 2e, Supplementary Figure 2i**). These studies, therefore, uncovered direct HNF1A-dependent genetic programs. They show that the function of HNF1A in pancreatic acinar cells entails direct transcriptional activation of a broad differentiated cell program that controls metabolic functions. They also revealed that HNF1A suppresses growth promoting pathways and that, given the absence of enriched HNF1A binding to upregulated genes, this is largely mediated through indirect regulatory mechanisms.

### Deregulation of the *ANFIA*-dependent program in non-classical PDAC

To explore the role of HNF1A-dependent programs in human PDAC, we asked if deregulated genes from *Hnf1a^aKO^* pancreas exhibit altered expression in human PDAC primary tumors. We thus examined transcriptome data from the TCGA-PAAD study ^35^. We defined human orthologs of genes that showed significant down or upregulation in *Hnf1a^aKO^* pancreas, and found that these gene sets showed concordant down or upregulation in *non-classical* tumor molecular subtypes – variously defined as quasimesenchymal, basal, and squamous-like PDAC ^3, 36, 37^ (**Figure 3a**). We also employed the same gene sets to cluster human tumor samples from TCGA-PAAD and ICGC-PACA cohorts ^3^, using non-negative matrix factorization. This exposed a cluster of predominantly non-classical tumors that showed transcriptional changes in the same direction as *Hnf1a^aKO^* pancreas (HNF1A cluster 3, **Supplementary Figure 3a**).

**Figure 3.**
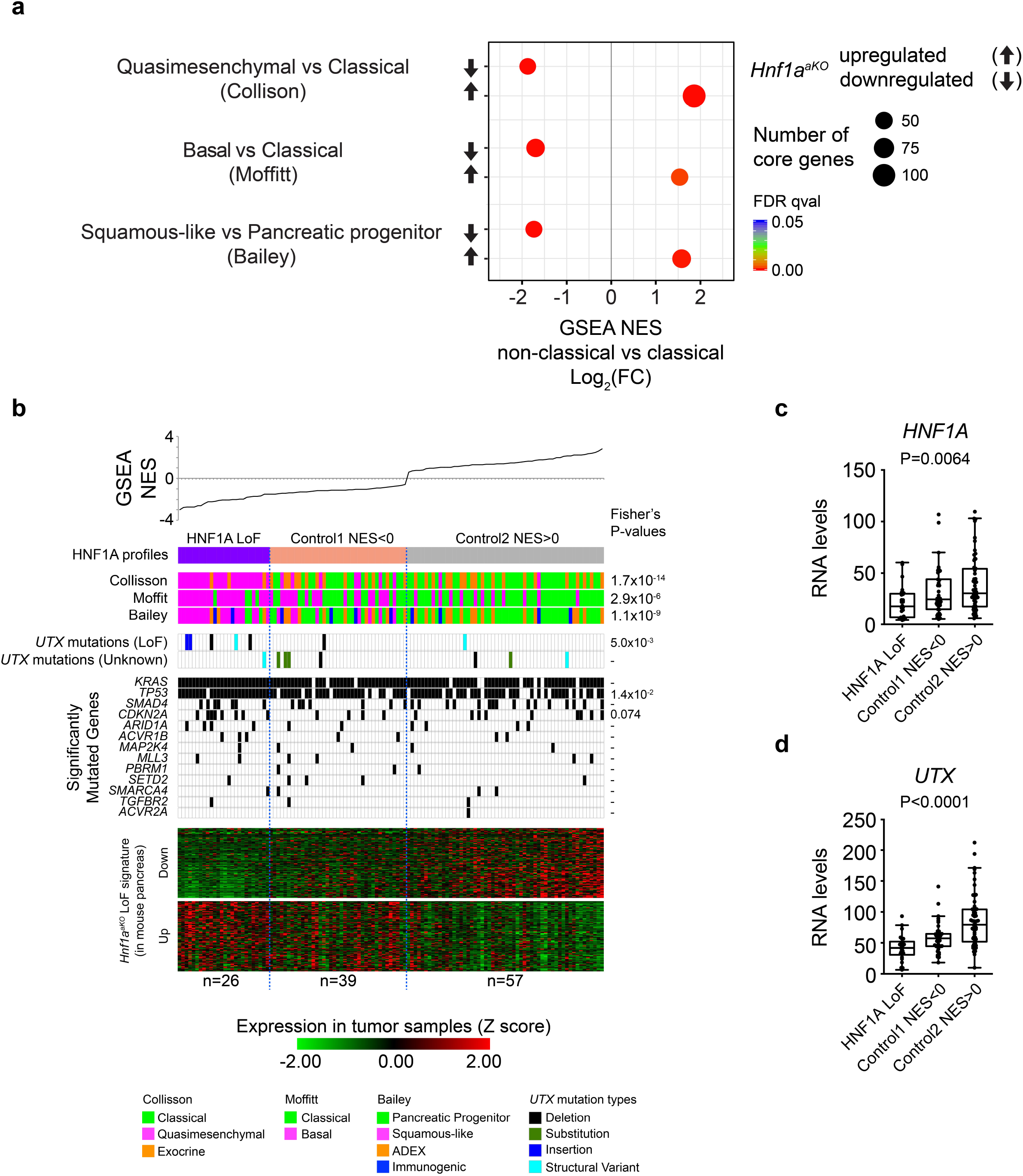
Human non-classical and UTX-deficient PDAC exhibit HNF1A-deficient programs. **a**, GSEA showed that down- or upregulated genes in *Hnf1a^aKO^* mice were down- or upregulated, respectively, in non-classical PDAC vs. classical molecular subtypes from the TCGA-PAAD study ^2^. The arrows denote down- or upregulated gene sets in *Hnf1a^aKO^* mice, the size of circles is proportional to the number of core-enriched genes in the GSEA analysis, NES denotes positive or negative normalized enrichment scores of gene sets in the human tumors, colors denote FDR q-values. **b**, Analysis HNF1A function in 121 high-purity cases of the ICGC-PACA-AU cohort identified tumors with most pronounced downregulation of direct HNF1A target genes. We performed GSEA with a gene set of 106 human orthologs of HNF1A direct targets showing downregulation in *Hnf1a^aKO^* pancreas. For each sample, GSEA of this gene set was performed against a gene list rank-ordered by differential expression to all other samples. Samples were ranked by the resulting normalized enrichment score (NES), and classified as either *Hnf1a* LoF samples (purple, NES < 0; P < 0.05), or *Control 1* (beige, NES < 0; P > 0.05) and *Control 2* (grey, NES>0). *Hnf1a* LoF samples are predominantly non-classical tumors according to Collisson, Moffitt and Bailey signatures ^3-5^. Putative loss of function *UTX* mutations (*UTX* LoF mutants) were found in 19% of *Hnf1a* LoF tumors vs. 2% of all others (Fisher’s P = 0.005). *UTX* mutants were considered functional if classified as high’ functional impact in ICGC (small <=200bp deletions, single base substitutions and insertions (small <=200bp), or if they were regarded as likely loss of function structural variants in Bailey et al. ^5^ All of these *UTX* mutation types were frame-shift mutations. Other *UTX* mutations were classified as unknown. **c**, *Hnf1a* mRNA levels differed in *Hnf1a* LoF and control groups (Kruskal-Wallis, P<0.01), despite considerable variability and overlap between groups. Heatmaps show Z-score normalized expression of human orthologs of significantly down- and up-regulated genes in *Hnf1a^aKO^* pancreas. We display genes showing differential expression across the 3 *Hnf1a* profiles (q<0.05, by SAM multiclass analysis; 85% of 106 downregulated human orthologs and 60% of genes of 146 upregulated human orthologs). **d**, *UTX* mRNA levels were downregulated in *Hnf1a* LoF tumors (Kruskal-Wallis, P<0.001).

We next sought to identify tumors with most pronounced HNF1A-deficient function. Because our findings showed that HNF1A primarily functions as a direct activator, we defined tumors with most pronounced downregulation of genes that were both HNF1A-bound and downregulated in *Hnf1a* mutant mice (hereafter referred to as HNF1A loss of function, or HNF1A LoF tumors). We found that HNF1A LoF tumors showed a remarkable concordance with tumors that were consistently classified as non-classical molecular subtypes in independent studies, and were therefore significantly enriched in quasimesenchymal (Fisher’s P = 1.7×10^−14^), basal (P = 2.9×10^−6^), and squamous-like (P = 1.1×10^−9^) subtypes (**Figure 3b**). They also had increased *TP63* mRNA, a marker of squamous-like PDAC ^3, 15, 38^ (**Supplementary Figure 3b**).

HNF1A LoF tumors – as well as non-classical PDAC – showed significantly lower *HNF1A* mRNA levels (Kruskall Wallis P = 0.0064), although there was considerable overlap with control tumors, suggesting that abnormal HNF1A function cannot be exclusively explained by differences in *HNF1A* expression (**Figure 3c, Supplementary Figure 3c**). Furthermore, although HNF1A LoF tumors had non-classical molecular signatures – which correlate with high histological grade – HNF1A immunoreactivity was not significantly lower in high histological grade (poorly differentiated) tumors in tissue microarrays of human PDAC (n=102) (**Supplementary Figure 3e**). These results reveal abnormal *HNF1A* function in a subset of non-classical PDAC tumors.

### *HNF1A*-dependent changes in UTX-deficient non-classical PDAC

Although a subset of non-classical PDAC samples exhibit a gene expression profile that resembles that of pancreatic *Hnf1a-mutant* mice, HNF1A mRNA levels were not invariably altered, and so far recurrent somatic *HNF1A* mutations have not been reported in PDAC. This raised the question of why non-classical PDAC shows abnormal HNF1A function. Non-classical (e.g. squamous-like) PDAC have been shown to express low *UTX* mRNA, and are enriched in *UTX* somatic genomic defects ^3, 15^. Our studies also showed decreased *UTX* mRNA in non-classical PDAC (P < 10^−3^) (**Supplementary Figure 3d**) and decreased UTX immunoreactivity in poorly differentiated tumors (P = 0.03, n=94) (**Supplementary Figure 3f**). Importantly, tumors with *HNF1A* LoF phenotypes showed decreased *UTX* mRNA (median [IQR]: 41.9 [32.2-50.5] in LoF tumors, vs. 57.3 [44.3-64.3] and 79.4 [53.1-104.2] in control tumors, Kruskall Wallis P < 10^−4^) (**Figure 3d**). Furthermore, the analysis of genotypes from the Australian ICGC-PACA study revealed putative loss of function *UTX* mutations in 19% of tumors showing *HNF1A* LoF phenotypes, compared with 2% of all other tumors (Fisher’s P = 0.005) (**Figure 3b**). Likewise, tumors with *UTX* putative loss of function mutations showed abnormal *HNF1A-dependent* programs (**Figure 3b**). Collectively, these correlations hinted at a mechanistic link between *UTX-* and *HNF1A*-deficient phenotypes in non-classical PDAC.

### *Utx*-dependent Kras-induced oncogenesis

To examine the relationship between *Hnf1a-* and *Utx*-deficiency in pancreatic cancer, we generated mice with pancreas-specific inactivating *Utx* mutations and oncogenic *Kras* mutations (*Pdx1^Cre^, Utx^LoxP/LoxP^*, *Kras^G12D^*, hereafter referred to as *Utx^pKO^*;*Kras^G12D^* mice; (**Supplementary Figure 4a-c**). We focused on female mice because UTX is encoded in the X chromosome, and males harbor a Y chromosome paralog named *Uty. Utx^pKO^*;*Kras^G12D^* mice showed normal pancreas morphology at 7 days of age (**Figure 4a-c**), but subsequently rapidly developed invasive PDAC. Early tumors were apparent by weaning, showing prominent signs of spindle cell proliferation, sarcomatoid morphology with occasional glandular tumor components, regions with widespread signs of ADM and abundant desmoplastic reaction (**Figure 4e-f and Figure 4h-i**). Subsequently, tumors showed massive remodeling and very rapid infiltrative growth (**Figure 4k-l and Figure 4n-o**), leading to the death of all *Utx^pKO^*;*Kras^G12D^* female mice by 15-16 weeks of age (**Supplementary Figure 4d**). This contrasted with 8 week-old control *Kras^G12D^* mice, which expectedly only presented occasional acinar-to-ductal metaplasia (**Figure 4j and m**). This finding was *consistent* with recently reported findings in 6 week-old mice ^15^. Male mice presented delayed mortality compared to females, suggesting partial *Uty* compensatory tumor suppressor functions, as reported previously ^15, 39^. These findings, therefore, showed *Utx*-dependent sarcomatoid PDAC lesions that are reminiscent of those observed in *Hnf1a^pKO^*;*Kras^G12D^* mice, although *Utx*-deficient tumors showed earlier age of onset and more rapid growth. These results were consistent with the overlapping *UTX*- and *HNF1A*-deficient molecular signatures of human tumors.

**Figure 4.**
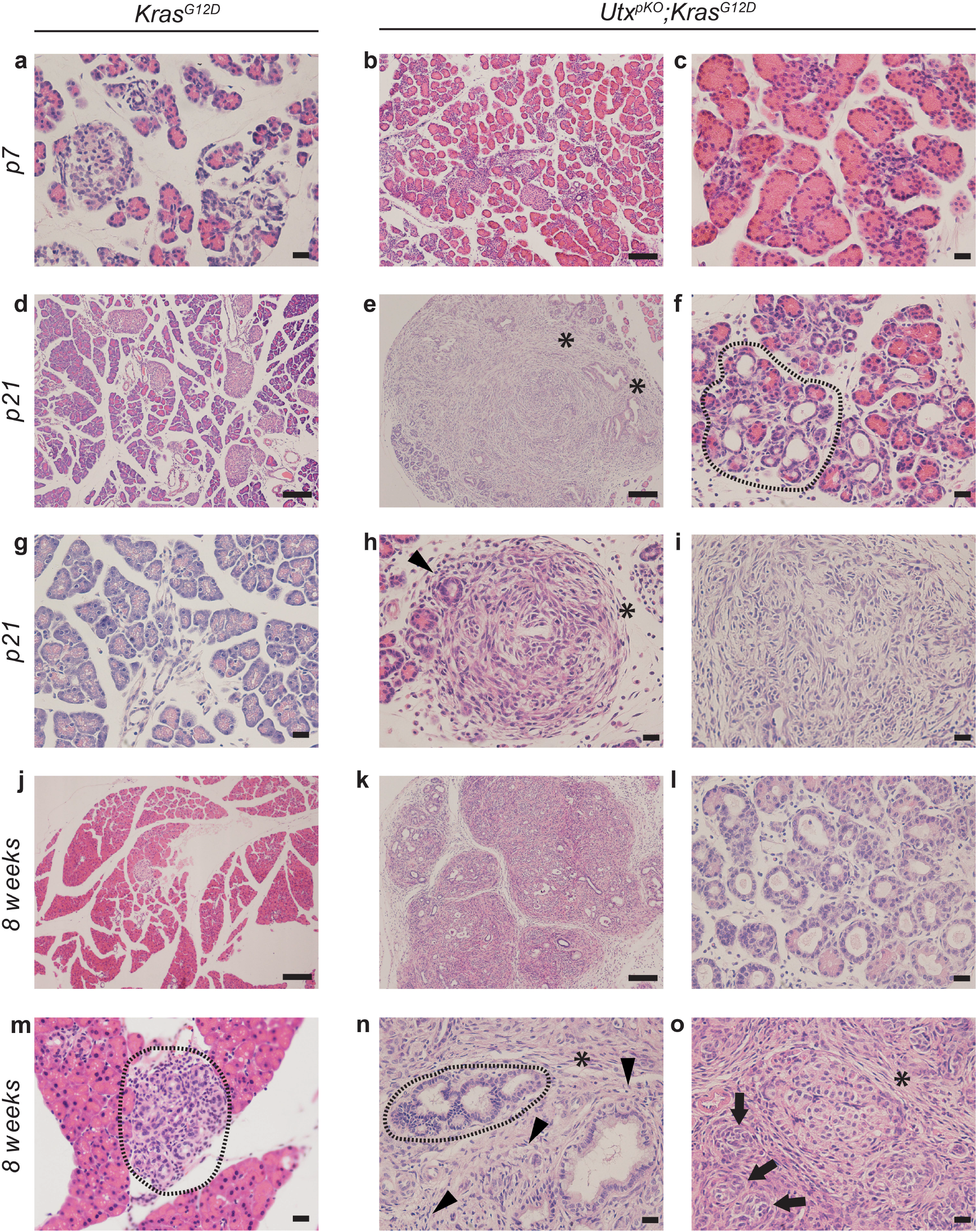
Utx-dependent Kras-induced oncogenesis. H&E staining of pancreata from *Utx^pKO^*;*Kras^G12D^* and *Kras^G12D^* mice. **a-c**, 7 day-old *Kras^G12D^* and *Utx^pKO^*;*Kras^G12D^* mice show normal morphology. **d-i**, 21 day-old, *Utx^pKO^;Kras^G12D^* mice already show acinar-to-ductal metaplasia (dashed encircled areas), spindle cell proliferation (asterisks), sarcomatoid architecture (i) and desmoplastic reaction (black arrowhead in h). **j-o**, *Kras^G12D^* mice present occasional acinar-to-ductal metaplasia and low grade PanINs at 8 weeks (j, m and see Figure 1n) whereas at the same age pancreas from *Utx^pKO^*;*Kras^G12D^* mice show massive remodeling (k), extensive acinar-to-ductal metaplasia (l and dashed encircled area in n), cancer with prominent spindle/mesenchymal proliferation, infiltrative growth (asterisk and black arrows in o) and abundant stroma (black arrowheads in n). Scale bars100 μm (b,d,e,j,k) or 20 μm (c,f,g-i, l-n).

### UTX activates a growth-suppressing program of differentiated acinar cells

The analysis of molecular signatures in human tumors, and the semblance of mouse genetic phenotypes, suggested that UTX and HNF1A might control common tumor suppressive programs in pancreatic exocrine cells. We thus studied the transcriptional function of UTX in non-tumoral, differentiated pancreatic exocrine cells, following a strategy similar to that described above for HNF1A. We examined female *Utx^pKO^* mice, in which the *Pdx1^Cre^* transgene led to pancreatic inactivation of *Utx* in most cells from all pancreatic epithelial lineages (>98% of which are exocrine cells) as early as e15.5 (**Supplementary Figure 5a**), and a marked reduction of UTX protein at weaning (**Supplementary Figure 4c**).

*Utx^pKO^* mice were born at the expected Mendelian ratio and appeared healthy. They had normal glycemia at 12 weeks of age (**Supplementary Figure 5b**) and unaltered pancreas morphology at birth and until weaning, indicating that, like HNF1A, UTX is dispensable for pancreas organogenesis (**Supplementary Figure 5c-f**). In analogy to observations in *Hnf1a^aKO^* mice, acinar cells showed increased proliferation at weaning (**Supplementary Figure 5i**). Unlike *Hnf1a^aKO^* mice, as *Utx^pKO^* female mice aged they showed acinar cell attrition, and by 8 weeks of age some atrophic pancreatic lobules could be observed (**Supplementary Figure 5g-h**).

To assess the transcriptional function of UTX in the pancreas, we analyzed pancreatic RNA from 4-day old female *Utx^pKO^* mice, prior to the development of any morphological acinar cell abnormalities. This revealed profound transcriptional changes, with downregulation of acinar-specific genes (but not duct- or islet-genes) and upregulation of mesenchymal genes (**Figure 5a, b, Supplementary Table 3**). Downregulated genes were associated with metabolic pathways including glutathione, peptide hormone, ether lipid metabolism, and NRF2 (antioxidant) metabolism (**Figure 5c**). Upregulated genes in *Utx^pKO^* mice were enriched in programs regulating cholesterol biosynthesis, extracellular matrix organization and the innate immune response (Fc gamma R-mediated phagocytosis). GSEA showed prominent enrichment in additional annotations related to oncogenic pathways, including EMT, Wnt, ErbB, MAPK, onctostatin M, and NFKB signaling (**Figure 5d, Supplementary Figure 5j, Supplementary Table 4**). The biological pathways that were down- and upregulated in *Utx^pKO^* mice were generally deregulated in the same direction as in *Hnf1a^aKO^* (**Supplementary Figure 5k, Supplementary Table 5**). These observations suggested that *Utx* could exert its tumor suppressor function in acinar cells through the regulation of broadly similar pathways as *Hnf1a*, namely through the maintenance of acinar differentiated cell programs and inhibition of growth-promoting pathways.

**Figure 5.**
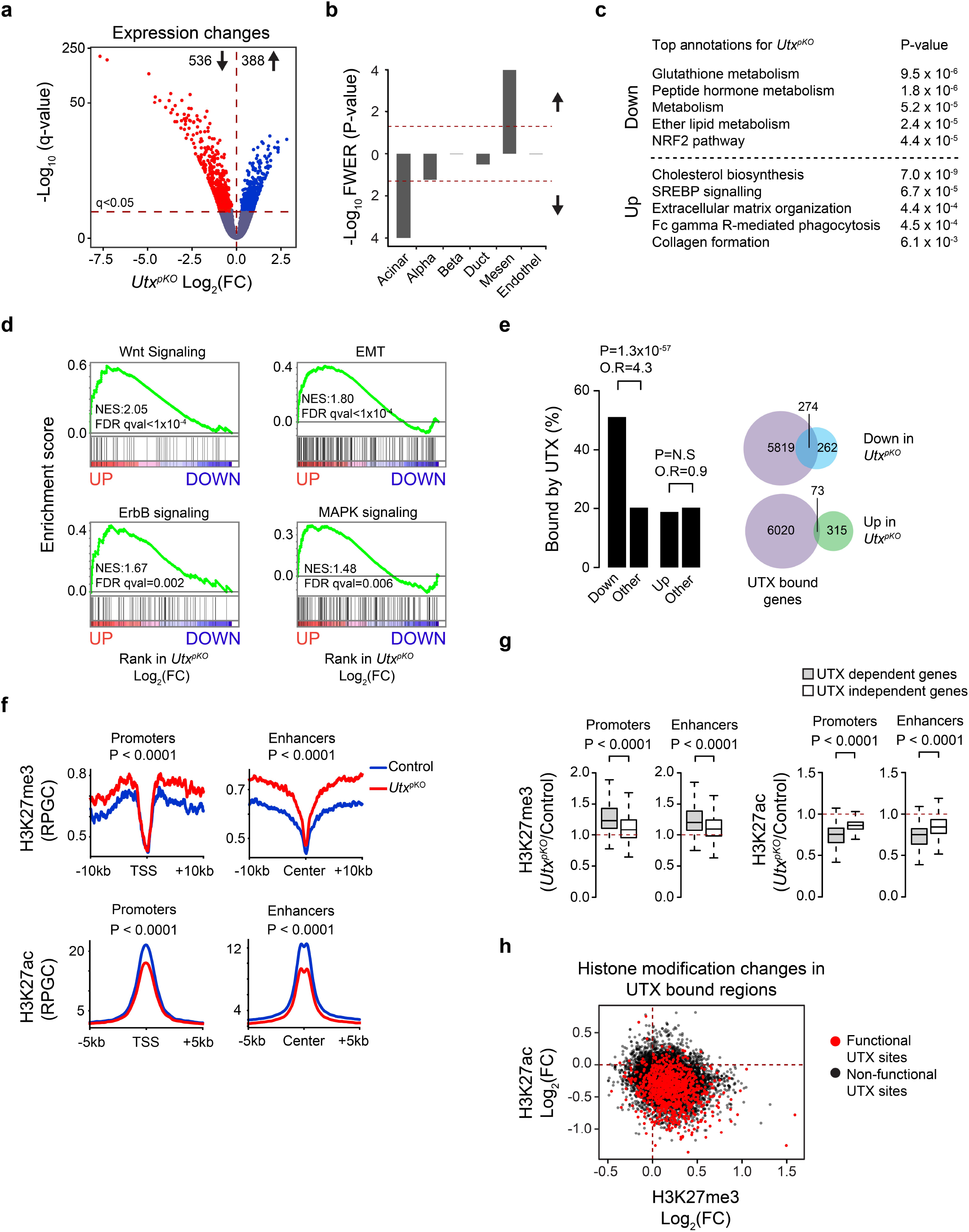
UTX promotes acinar cell differentiation gene programs and suppresses oncogenic pathways. **a**, Fold change (FC) in transcripts in *Utx^pKO^* vs. control pancreas, plotted against significance (-Log_10_ q; genes significant at q<0.05 are shown as colored dots above the horizontal line). **b**, GSEA showing that genes specific to differentiated acinar cells were downregulated in *UTX-deficient* pancreas, but not genes specific to islets or duct cells. Upregulated genes were enriched in genes specific to mesenchymal cells. Lineage-enriched genes were obtained from Muraro et al. ^1^ **c**, Functional annotation of most enriched processes in genes differentially expressed in *Utx^pKO^.* **d**, GSEA showed increased expression of indicated oncogenic pathway genes in *Utx^pKO^* pancreas. **e**. UTX functions as a transcriptional activator of direct target genes in pancreatic cells. Left: Downregulated, but not upregulated, genes in *Utx^pKO^* pancreas were enriched for UTX binding. P-values and odds ratios (O.R.) were calculated by Fisher’s exact test. Right: Venn diagrams showing overlap of down and upregulated genes with UTX binding. **f**, Aggregation plots of H3K27me3 and H3K27ac signals in UTX-bound enhancers and promoters show increased H3K27me3 and decreased H3K27ac in *Utx^pKO^* relative to control pancreas. **g,h**, UTX-bound genes that showed decreased expression in *Utx^pKO^* pancreas (*functional* UTX binding sites) showed simultaneous gain of H3K27me3 and loss and of H3K27ac. P-values in (**g**) were determined by two-tailed Mann-Whitney U test.

### Shared transcriptomes in *UtX^pKO^* mice and *UTX*-mutant human PDAC

We assessed the relevance of the transcriptional phenotype of *Utx^pKO^* pancreas to human PDAC. GSEA showed that up- and downregulated genes from tumors with *UTX* putative loss of function mutations were concordantly up- and downregulated in *Utx^pKO^* pancreas (**Supplementary Figure 5l**). Furthermore, low *UTX* mRNA was observed in human tumors that showed greatest deregulation of genes downregulated in *Utx^pKO^* pancreas (**Supplementary Figure5m**). Likewise, non-classical PDAC tumors, which exhibit decreased *UTX* expression, showed down- and up-regulation of deregulated genes in *Utx^pKO^* pancreas (FDR q = 10^−4^ and 0.04, respectively; **Supplementary Figure 5n**). The transcriptional changes observed in *Utx^pKO^* mice are, therefore, directly relevant to a subset of human PDAC tumors.

### UTX co-activates acinar differentiated cell programs

To further understand the direct mechanisms whereby UTX regulates genetic programs relevant to PDAC, and the potential link to HNF1A, we integrated transcriptome changes in *Utx^pKO^* pancreas with profiling of UTX-bound genomic sites in pancreas from 4-day old mice. We identified 8455 UTX binding sites (**Supplementary Figure 5o**), which were not detected when performing ChIP-seq in *Utx^pKO^* pancreas (**Supplementary Figure 5p**). Most UTX binding sites were located in active enhancers and promoters (**Supplementary Figure 5q**), and they were enriched in genes that were downregulated in *Utx^pKO^* mice, but not in those that showed upregulation (**Figure 5e, Supplementary Figures 5p and 2g**). UTX was, consequently, preferentially bound near acinar-enriched genes, but not to genes involved in EMT or other pathways that showed induction in *Utx^pKO^* pancreas. This indicates that although UTX has been reported to have transcriptional activating and repressive functions ^39^, the most prevalent direct function of UTX in pancreatic cells appears to be the transcriptional activation of gene targets, and indirect inhibition of many other genes.

UTX is a H3K27 demethylase, although previous studies have also demonstrated UTX functions that are independent from this catalytic activity ^15, 39–43^. We observed that in the pancreas of *Utx^pKO^* mice, UTX-bound enhancer and promoter regions showed increased H3K27me3 (**Figure 5f**), which was greatest in genes that showed significant downregulation in *Utx^pKO^* mice (**Figure 5g, Supplementary Figure 5p**). Consistent with UTX forming part of complexes containing histone acetyl transferases p300/CBP ^44–46^ we also observed decreased H3K27ac in those regions in *Utx* mutants (**Figure 5f,g Supplementary Figure 5p**). UTX-bound sites, therefore, often showed concomitant H3K27me3 gain and H3K27ac loss (**Figure 5h**). Collectively, these results indicate that the direct function of UTX in pancreatic cells entails profound chromatin changes and transcriptional activation of genomic binding sites.

### HNF1A and UTX share functional targets in acinar cells

UTX is not a sequence-specific DNA binding protein and the mechanism that recruits UTX to its genomic sites is poorly understood. We therefore examined UTX-bound sequences to identify candidate transcription factors (TFs) that recruit UTX in pancreatic cells, and specifically focused on the subset of UTX-bound sites associated with transcriptional changes in *Utx^pKO^* mice. These *functional* UTX-bound regions were enriched in *in silico*-predicted DNA-recognition sequences for canonical acinar TFs such as FOXA, GATA6, NR5A2 and RBPJL, although the most enriched sequence was the HNF1 recognition motif (**Figure 6a**). Consistently, functional UTX-bound regions were enriched for *in vivo* binding sites for multiple acinar cell TFs, yet showed the highest enrichment for HNF1A binding (**Figure 6b**). These findings, therefore, pointed to a strong overlap between UTX and HNF1A occupancy in pancreatic cells. Consistent with these findings, 31 of the 40 most downregulated genes in *Utx^pKO^* pancreas were also downregulated in *Hnf1a^aKO^* pancreas, or are known to be HNF1A-depdendent target genes (**Figure 6c**). Accordingly, gene set enrichment analysis (GSEA) showed that genes that were significantly down- or upregulated in *Utx^pKO^* mice were preferentially down- or upregulated in *Hnf1a^aKO^* mice (**Figure 6d**, and **Figure 6e** for the converse comparison). This indicated that the derangement of shared biological pathways in *Utx^pKO^* and *Hnf1a^aKO^* mice (shown in **Supplementary Figure 5k**) was largely due to deregulation of common genes. More specifically, UTX-bound genes that showed downregulation in *Utx^pKO^* mice were largely downregulated in the pancreas of *Hnf1a^aKO^*, (**Figure 6f and 6g**). Importantly, UTX-bound genes showed downregulation in *Utx^pKO^* if they were co-bound by HNF1A, but showed more marginal changes when they were not co-bound by HNF1A, suggesting that HNF1A could be critical for UTX function (**Figure 6h**). Collectively, these findings indicate that HNF1A and UTX target common genomic sites and regulate shared gene programs in pancreatic acinar cells.

**Figure 6.**
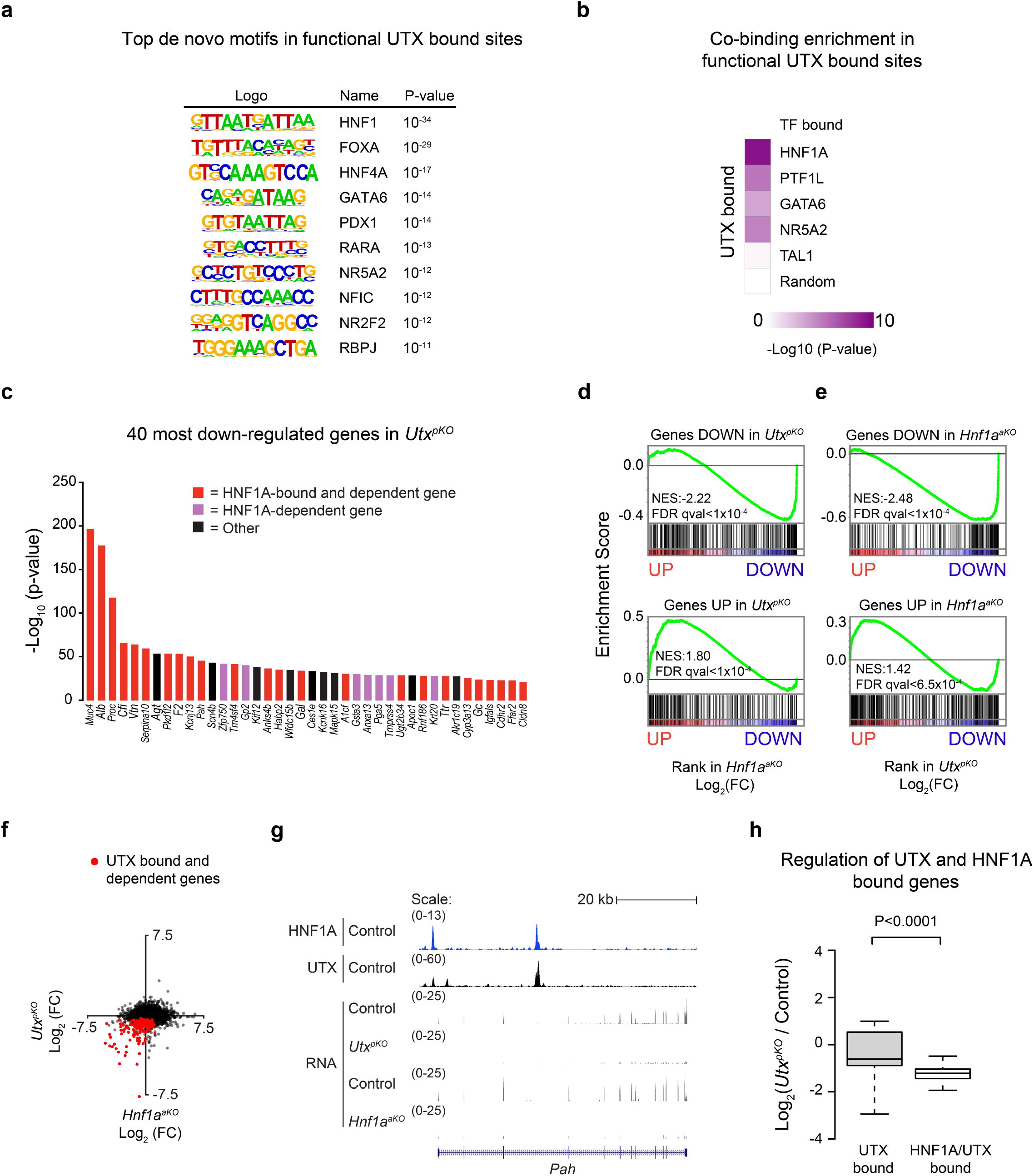
UTX and HNF1A regulate shared target genes. **a**, Motif analysis in functional UTX-bound regions, showing top ten *de novo* motifs ranked by P-value determined by HOMER software. **b**, Co-binding analysis in functional UTX-bound enhancer and promoter regions revealed that HNF1A was the most enriched co-bound TF amongst three other acinar cell TFs. Binding regions of TAL1 in a non-pancreatic cell type and random binding sites were used as negative controls. P-values was determined by Fisher’s exact test for peak comparisons using all enhancer and promoter regions as background. **c**, The most downregulated genes in *Utx^pKO^* pancreas are shown ranked by q-value, and are almost invariably bound by HNF1A and down regulated in *Hnf1a^aKO^* pancreas, or known to be direct HNF1A-dependent target genes from other studies (red and purple, respectively). **d,e**, GSEA analysis on the *Hnf1a^aKO^* and *Utx^pKO^* ranked-ordered gene lists vs. their reciprocal up- or downregulated gene sets, demonstrated that UTX and HNF1A regulate similar genes. **f**, Scatter plot showing expression changes of *Hnf1a^aKO^* and *UtX^pKO^* genes. Genes that were bound by UTX and downregulated in *Utx^pKO^* pancreas (functional UTX targets; red dots) were downregulated in *Hnf1a^aKO^* pancreas. **g**, Example showing that HNF1A and UTX co-occupy the same regions in the *Pah* gene. In *Hnf1a* or *Utx* knock-out pancreas, *Pah* expression is downregulated. **h**, Genes that were co-bound by UTX and HNF1A showed greatest downregulation in *UtX^pKO^* pancreas as compared with UTX bound genes that were not bound by HNF1A depicted as box plots with median and interquartile range of Log_2_ fold-changes in TPM values. P-values were determined by two-tailed Student’s t tests.

### HNF1A recruits UTX to genomic targets in pancreatic cells

To define the molecular mechanisms that link UTX and HNF1A function, we performed co-immunoprecipitation experiments using purified nuclei from a mouse acinar cell line. This showed that HNF1A and UTX form part of a common complex in pancreatic cells (**Figure 7a**). To test whether this interaction could mediate the recruitment of UTX by HNF1A to its genomic targets, we performed ChIP-seq for UTX using chromatin from pancreas of wild type and *Hnf1a^-/-^* mice. This showed that UTX binding to genomic targets was drastically reduced in *Hnf1a*^-/-^ pancreas (**Figure 7b**). Importantly, regions with reduced UTX binding in *Hnf1a*^-/-^ pancreas were bound by HNF1A and carried HNF1 motifs (**Figure 7c-e**). These findings, therefore, showed that HNF1A recruits UTX to genomic targets. Consistent with the functional importance of this recruitment, most genes associated with reduced UTX binding in *Hnf1a*-deficient pancreas were direct HNF1A target genes with decreased expression in *Hnf1a*-deficient pancreas (**Figure 7f, Supplementary Figure 6a-e**). By contrast, HNF1A binding was not affected in *Utx^pKO^* pancreas, indicating that, despite the functional interdependence of both proteins, UTX is not required for HNF1A binding to chromatin (**Supplementary Figure 6f**). These results, therefore, show that HNF1A interacts in a common complex with UTX, and recruits it to its functional targets in pancreatic cells. Collectively, these findings suggest a mechanistic model (**Figure 7g**) that explains the overlapping genomic phenotypes of HNF1A- and UTX-deficiency in genetic mouse models and human tumors, and their contribution to determine PDAC subtypes.

**Figure 7.**
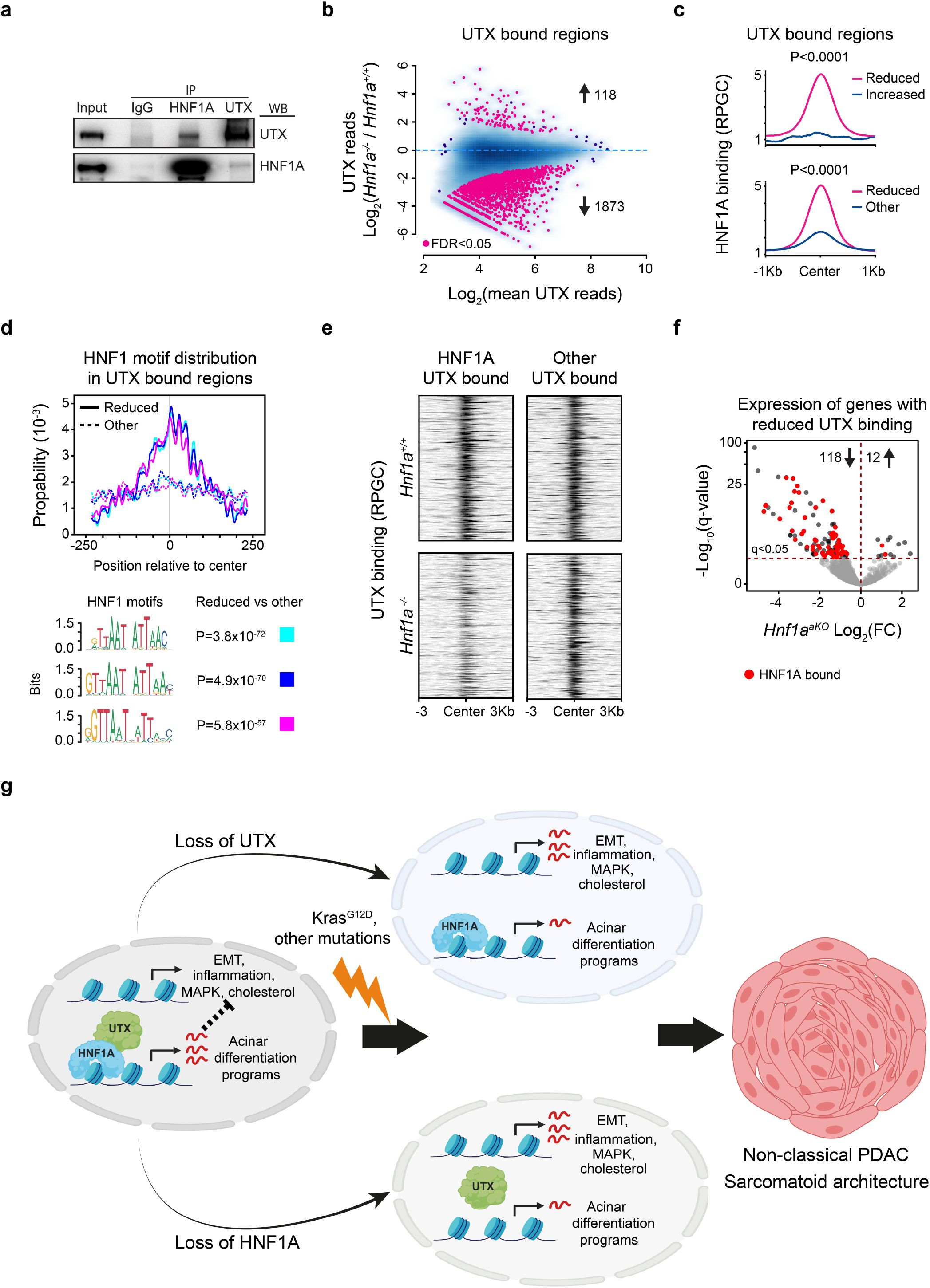
HNF1A recruits UTX to activate transcription of its target genes. **a**, Co-immunoprecipitation experiment of endogenous HNF1A and UTX followed by western blot demonstrated that HNF1A is in the same complex as UTX. **b**, Differential binding analysis of UTX in *Hnf1a*^-/-^ vs. wild type pancreas. Pink dots below zero (1873 sites) show regions with reduced UTX binding and pink dots above zero (118 sites) are regions with increased binding at FDR<0.05. **c,d**, Regions that show reduced Utx binding in *Hnf1a*^-/-^ chromatin are strongly bound by HNF1A and are highly enriched in HNF1 motifs. P-values in (**c**) were calculated with two-tailed Mann-Whitney U test and in (**d**) with Fisher’s exact test. **e**, UTX binding is markedly reduced in HNF1A and UTX co-bound regions in *Hnf1a^-/-^* pancreas, but not in other UTX bound regions. **f**, Genes that loose UTX binding in Hnf1a-mutant pancreas are predominantly downregulated in *Hnf1a^aKO^* pancreas and are direct HNF1A target genes (red dots). **g**, Summary model depicting that HNF1A recruits UTX to genomic binding sites to activate an acinar differentiation program that indirectly suppresses core oncogenic pathways. In the presence of *KRAS* mutations and other oncogenic events, defective HNF1A or UTX function results in failure of this shared program, with increased activity of pathways that promote high-grade non-classical PDAC.

## Discussion

In this study we provide a direct genetic demonstration that *HNF1A*, which encodes for a homeodomain transcription factor best known for its causal role in autosomal dominant diabetes ^47^, has a tumor suppressive function in the exocrine pancreas. We further provide epigenomic, biochemical and genetic evidence that mechanistically link *HNF1A* function in pancreatic exocrine cells to *UTX*, an established tumor suppressor. These results in turn indicate that the tumor suppressive function of *UTX* is tightly linked to its role as a transcriptional co-regulator of epithelial differentiation. We demonstrate that HNF1A-deficiency partially phenocopies UTX-driven tumorigenesis in mice, and show that *HNF1A/UTX*-deficiency is a prominent feature of the genetic programs of non-classical human PDAC tumors. These findings, therefore, shed light on mechanistic underpinnings of emerging sub-classification of PDAC subtypes.

Our analysis of *Utr*-deficient *Kras^G12D^* mice is consistent with a recent report showing aggressive PDAC in *Utr*-deficient *Kras* mutant mice ^15^, and extend those studies by defining the genetic programs that are directly regulated by UTX in the differentiated pancreatic exocrine cells that give rise to PDAC. The UTX-deficient transcriptome phenotype observed in non-tumoral pancreas was consistent with that reported in tumors from *Utr*-deficient *Kras* mutant mice ^15^. Our integrated analysis of genomic binding sites indicated that the positive regulatory effect of UTX (and HNF1A) on acinar differentiation genes are largely direct, whereas the suppression of oncogenic and EMT regulatory programs in pancreatic cells is predominantly indirect.

Our studies, therefore, exposed a molecular mechanism whereby HNF1A recruits UTX to activate differentiation programs of pancreatic exocrine epithelial cells, thereby suppressing oncogenic and EMT pathways (**Figure 7g**). This mechanism aligns well with the observation that *Utx-* and *Hnf1a*-deficient tumors in mice featured sarcomatoid architecture. We further showed that *HNF1A-* and *UTX-* dependent pancreatic programs are defective in non-classical PDAC tumors, variously defined as human basal, quasi-mesenchymal or squamous molecular types ^3, 35–37, 48^. Interestingly, abnormalities in the *HNF1A/UTX* dependent pancreatic program in tumors were associated with either somatic *UTX* mutations, or to decreased expression of *UTX* or *HNF1A*, in consonance with the proposed molecular mechanism.

These findings indicate that the disruption of *HNF1A* and *UTX*-dependent programs not only promotes the formation of PDAC, but is also instructive for the definition of PDAC subtypes. *HNF1A* is expressed in normal acinar cells, the presumed cell of origin of most PanIN precursors and PDAC ^33, 34^, but not in duct cells. The results suggest that in the presence of other oncogenic events, defective *HNF1A*-, *UTX*-dependent gene regulation can promote the development of an undifferentiated high-grade PDAC cellular phenotype (**Figure 7g**). More generally, our results open avenues to study the precise manner in which *HNF1A-* and *UTX*-dependent regulation of epithelial differentiation programs fails and thereby contributes to the development of PDAC.

## Acknowledgements

This research was supported by the National Institute for Health Research (NIHR) Imperial Biomedical Research Centre. Work was funded by grants from the Wellcome Trust (WT101033 to J.F.), Medical Research Council (MR/L02036X/1 to J.F.), European Research Council Advanced Grant (789055 to J.F.), Ministerio de Economía y Competitividad (BFU2014-54284-R, RTI2018-095666-B-I00 to J.F., SAF2011-29530 and SAF2015-70553-R to F.X.R. and RTICC from Instituto de Salud Carlos III (RD12/0036/0034, RD12/0036/0050) to F.X.R. M.K. was supported by a Juvenile Diabetes Research Foundation postdoctoral fellowship (3-PDF-2014-192-A-N). I.M. was supported by a Fellowship from Fundació Bancaria La Caixa (ID 100010434) (grant number LCF/BQ/ES18/11670009). Work in CRG was supported by the CERCA Programme, Generalitat de Catalunya. Work at CRG and CNIO were supported by Ministerio de Ciencia, Innovación y Universidades as Centro de Excelencia Severo Ochoa (SEV-2012-0208, SEV-2015-0510). We thank the University of Barcelona School of Medicine animal facility, Center of Genomic Regulation and Imperial College London Genomics Units, and the Imperial College High Performance Computing Service. We thank I. Cobo, S. Paliwal and members of the Ferrer lab for valuable discussions.

## Materials and Methods

### Animal Studies

Animal experimentation was carried out in compliance with the EU Directive 86/609/EEC and Recommendation 2007/526/EC regarding the protection of animals used for experimental and other scientific purposes, enacted under Spanish law 1201/2005.

The *Utx^LoxP^* ^1^, *Pdx1^Cre^* ^2^, *LSL-Kras^G12D^* ^3^, *Hnf1a*^-/-4^, and *Ptf1a^Cre^* ^5^ mouse lines have been described.

To generate a conditional *Hnf1a* allele, a floxed *Hnf1a* exon 2 was generated in C57Bl/6N JM8.F6 embryonic stem cells (ES cells) ^6^ by homologous recombination. Briefly, ES cells were electroporated with a targeting plasmid construct linearized by PmeI and MluI, containing the floxed exon and a PGK/Neomycin selection cassette flanked by FRT recombination sites, and then selected with Geneticin. Several correctly targeted ES cell clones were identified by Southern blot using external probes detecting a 9.2kb 5’ ScaI fragment, an 8.7kb 3’ BamHI fragment from the recombined allele and a neo probe detecting a unique 11.7 kb SpeI Neomycin fragment indicating no additional construct integrations. The presence of the distal LoxP site, upstream of exon 2, was confirmed by PCR using the primers indicated in **Supplementary Table 6**. Correctly targeted ES cell clones were injected into C57BL/6BrdCrHsd-Tyrc morulae (E2.5) to create chimeric mice that transmitted the recombined allele through the germ line. The PGK/Neo cassette was excised by crossing heterozygous mice with Tg.CAG-F1p mice ^7^, which express the FLPe recombinase in the germ line to create the *Hnf1a* conditional knockout allele. Lines with floxed alleles without Cre, Cre lines without floxed alleles, *Pdx1^Cre^*;*LSL-KRAS^G12D^* or wild-type littermates served as controls. Gender was considered as a variable and included in the study design. Genotype was confirmed by PCR using the primers provided in **Supplementary Table 6**.

Prior to decapitation, mice were anesthetized using isoflurane (Zoetis), pancreas was collected quickly and placed in paraformaldehyde 4% (Sigma-Aldrich) for paraffin embedding or processed for RNA isolation.

### Histology, TMA preparation and Immunohistochemistry

Fixation of mouse pancreas was performed using paraformaldehyde 4% (Sigma-Aldrich) or 4% buffered formalin overnight at 4°C followed by dehydration using increasing concentrations of ethanol. Dehydrated samples were embedded in paraffin blocks and cut in 4 μm sections. Sections were then deparaffinized and rehydrated. For hematoxylin and eosin analysis, the pancreata were stained, dehydrated, mounted with DPX (Panreac) and photo documented using optical microscopy (Olympus BX41TF).

Tissue samples for tissue micro arrays (TMAs) were obtained from patients who underwent pancreatic resection for PDAC (n=223) at the Department of Surgery of the University Hospital of Duesseldorf, Germany. The samples were fixed in 4% formalin and subsequently routinely embedded in paraffin. Three tumor samples per case (two samples from the tumor center, one from the periphery) with a 1-mm core size were selected and assembled into the TMA (Manual Tissue Arrayer MTA-1, Beecher Instruments, Inc., Sun Prairie, WI, USA). The use of human tissue samples was approved by the local ethics committee at the University Hospital of Duesseldorf, Germany (study number 5387). The TMAs were then cut in 2 μm sections for subsequent procedures.

Immunohistochemistry on mouse pancreas or TMAs, was performed according to manufacturer’s instructions using VectaStain^®^ ABC HRP Rabbit IgG kit (Vector Laboratories, Peterborough, UK). Epitope retrieval was achieved by pretreatment with boiling citrate buffer (10 mM citric acid, pH6) for 15 min using a pressure cooker. Endogenous peroxidase and protein blocking was performed with 3% H_2_O_2_ diluted in PBS for 10 min and with 1% BSA, 10% normal goat serum (Abcam, Cambridge, UK) and 0.1% Triton-X (Merck KGaA, Darmstadt, Germany) for 60 min. Anti-HNF1A and anti-UTX stainings were performed at a dilution of 1:250 (Anti-HNF1A, Abcam ab204306, Cambridge, UK), 1:200 (Anti-HNF1A, Cell Signaling Technology, 89670, Leiden, The Netherlands) and 1:100 (Anti-UTX, Cell Signaling Technology 33510S, Denver, USA), respectively. Diaminobenzidine (Abcam ab64238, Cambridge, UK) was used as chromogen to visualize protein expression, counterstaining was achieved with Mayer’s hemalaun solution (Merck KGaA, Darmstadt, Germany).

### Analysis of TMAs

The semi-quantitative analysis of the stained sections was done by light-microscopy according to the immunoreactive score (IRS) of Remmele and Stegner ^8^. Both the staining intensity and the number (%) of positive stained cells were evaluated for each case. If more than one tumor core was available in one case, a mean value was calculated. Staining intensity was scored as no staining (0), weak staining (1), moderate staining (2) or strong staining (3). The proportion of stained cells was scored as follows: no staining=0, 1-9%=1, 10-50%=2, 50-80%=3, >80%=4. The IRS was calculated by multiplication of the intensity value and the percentage value, resulting in a score between 0 and 12. Finally, IRS results were classified into four groups as follows: 0-1=negative (group 0), 2-3=weak (group 1), 4-8=moderate (group 2), 9-12=strong (group 3). Each tumor case was graded according to UICC and WHO classifications. Two cases with undetermined grade (Gx) were omitted from the analysis. The rest of the cases were either scored as G2 or G3 grades. Contingency table analysis and Chi-square test were used to evaluate the association between IRS classifications and tumor grades.

### Immunofluorescence

Immunofluorescence analysis of murine pancreata was performed on paraformaldehyde-fixed paraffin-embedded tissue cut into 4 μm sections, as described with modifications^9^. Antigen unmasking was performed boiling sections on 10 mM citrate buffer pH 6 for 10min and tissue was permeabilized with PBS 0.5% Triton X-100 (Sigma). Blocking was performed by 1h incubation in humid chamber with 3% of normal donkey serum (Vendor) in blocking solution (DAKO). Slides were incubated in humid chamber with primary antibodies overnight at 4°C and washed in PBS 0.2% Triton-X100. Slides were incubated with secondary antibodies for 1h in humid chamber at room temperature, washed for 5 min in PBS 0.2% Triton-X100 and mounted using fluorescence mounting medium (DAKO). Stained slides were analyzed using a confocal laser scanning microscope (Leica CTR6500). Primary antibodies used are indicated in **Supplementary Table 7**.

### Western blotting and co-immunoprecipitation

All protein isolation steps were performed at 4 °C or on ice and protein concentrations were assessed with the Bradford assay (Bio-Rad). Protein from total pancreas lysates was isolated from a small fragment of mouse pancreas and minced in lysis buffer (20 mM Tris-HCl, pH 7.5, 150 mM NaCl, 1 mM Na_2_EDTA, 1 mM EGTA, 1% Triton X-100, 2.5 mM sodium pyrophosphate, 1 mM b-glycerophosphate, 1 mM Na_3_VO_4_, 1 μg/ml leupeptin containing 1x phosphatase inhibitor cocktail, Sigma-Aldrich and 4 x EDTA containing complete protease inhibitor cocktail, Roche). Lysates were freeze-thawed twice, cleared at 15000 g for 15 min and then the supernatant was recovered. For Western blots, lysate supernatants were denatured in 2X NuPAGE LDS sample buffer (Thermo Fisher) containing 0.1 M DTT at 95 °C for 5 min. Equal amount of lysates were loaded in NuPAGE 4-12% Bis-Tris Protein Gels (Life Technologies) and run at 150v for 90min. Gels were transferred to 0.2μm pore PVDF membrane (Life Technologies) and blocked with 5% fat-free milk powder for 60 min and probed overnight (ON) with primary antibody. Horseradish peroxidase conjugated secondary antibodies were used at 1:10,000 in BSA 5% and incubated 1h at RT. The blot was developed using ECL detection reagent (Amersham ECL, GE Healthcare). The pancreatic acinar cell line 266-6 (ATCC CRL-2151) was grown in DMEM (Thermo Fisher) with 10 % fetal calf serum, 1 % pen/strep and passaged twice per week. For co-immunoprecipitation, the cells were washed in ice-cold PBS and resuspended in hypotonic lysis buffer (10 mM HEPES, pH 7.9, with 1.5 mM MgCl_2_ and 10 mM KCl). After plasma membrane disruption, the lysate was centrifuged at 4000g for 5min and the nuclei lysed for 20min with gentle shaking in nuclear extraction buffer (20 mM HEPES, pH 7.9, with 1.5 mM MgCl_2_, 0.42 M NaCl, 0.2 mM EDTA, and 25% (v/v) Glycerol) before removing cell debris by centrifugation at 15,000g for 15 min. The cleared nuclear lysate was equilibrated to 150 mM NaCl, pre-cleared with Dynabeads Protein G (Thermo Fisher) for 1 h at 4 °C with rotation and 4 mg of extract was incubated overnight with 2 μg of anti-Utx, anti-HNF1A or normal rabbit IgG. Immune complexes were precipitated with 20 μl Dynabeads for 2 h and washed four times with IP wash buffer (10 mM Tris-HCl, pH 8, 150 mM NaCl, 0.1% NP-40 and 1 mM EDTA) supplemented with protease inhibitor. Proteins were eluted with 2X NuPAGE LDS sample buffer (Thermo Fisher) containing 0.1 M DTT at 95 °C for 5 min, and analyzed by western blotting.

### RNA isolation

For gene expression profiling we implemented a method for purification of intact pancreatic RNA according to a modified Guanidinium salts method ^10^. Briefly, a small piece of pancreatic tissue was immediately homogenized with a Polytron (VWR) in precooled 4 M Guanidinium Thiocyanate with 112 mM beta-mercaptoethanol, centrifuged at 5000g for 5 min at 4°C to remove insoluble debris. RNA was precipitated from the supernatant with pre-cooled 75% Ethanol, 0.1M potassium acetate, pH 5.5, and 75mM acetic acid at −20 °C for 2 hr. The precipitate was pelleted by centrifugation at 10,000g for 10 min at 4 °C and resuspended at room temperature in 7.5 M Guanidinium HCl and 10.5 mM beta-mercaptoethanol. The RNA was re-precipitated twice with 0.1 M potassium acetate, pH 5.5, and 50% ethanol to remove residual RNAses, followed by purification with TRI Reagent RNA Isolation Reagent (Sigma-Aldrich). The concentration and quality of RNA was measured with a NanoDrop spectrophotometer (ND-1000, Thermo Scientific) and an Agilent 2100 Bioanalyzer. RNA integrity numbers ranged from 7.8 to 9.3.

Alternatively, we used a previously described protocol ^11^.

### RNA-seq

One μg of total RNA was used to make RNA-seq libraries using the Truseq Stranded mRNA Sample Prep Kit (Illumina) following manufacturer’s instructions and sequenced by an Illumina HiSeq2500 platform with single-end reads of 50 bases. Conversion to FASTQ read format was done using Illumina’s bc12fastq algorithm. Four or three pancreases from, respectively, 4 days old or adult female mice of each genotype were analyzed: *Utx^LoxP/LoxP^* and *Pdx1^Cre^*;*Utx^LoxP/LoxP^* or *Ptf1a^Cre^* and *Ptf1a^Cre^*;*Hnf1a^Loxp/Loxp^*. Raw RNA-seq reads were aligned to the mouse transcriptome (a combined build of cDNA and ncRNA from Mus musculus v.GRCm38.p5, release 87) and quantified using Salmon v0.7.2 12, with the following parameters: salmon quant --gcBias, -- libType A and --fldMean and -- fldSD. The latter two were set to the average size and standard deviation of the fragment length distribution of the given library, respectively. The counts were used in DEseq2 (v1.14.1) ^13^ with R (v3.3.3) ^14^ for normalization and identification of significant differential expression (q<0.05) between controls (n=4) and mutants (n=4). Only genes with log2 (basemean) > 2.5 were retained for downstream analysis. For visualization, STAR (v2.3.0) ^15^ was used to align reads to the mouse genome assembly GRCm38.p5 (mm10) and resulting BAM files were converted to bigwig with reads per genomic content (RPGC) normalization by Deeptools (v2.4.2) ^16^.

### ChIP sequencing

ChIP experiments were performed on pancreas from day 4 old or 3 weeks old female mice. A Polytron was used to quickly mince 1-2 dissected pancreas in PBS with protease inhibitors followed by crosslinking with 1% formaldehyde for 10min at room temperature on a rotator and then quenched with 0.125M glycine for 5min. Tissue pieces were washed in PBS and lysed in 130 μl lysis buffer (2% Triton X-100, 1% SDS, 100 mM NaCl, 10 mM Tris-HCl, pH 8.0, 1 mM EDTA) with protease inhibitor on ice for 10 min and finally resuspended with a 30G needle syringe. The chromatin preparation was then sonicated to fragments enriched in the size range of 150 to 500 bp using S220 Focused Ultrasonicator (Covaris) and centrifuged at 14000 rpm for 10 min at 4 °C to pellet insoluble material. The supernatant was incubated for 1 h with 350 μl RIPA-LS buffer (10 mM Tris-HCl, pH 8.0, 140 mM NaCl, 1mM EDTA, pH 8.0, 0.1% SDS, 0.1% Na-Deoxycholate, 1% Triton X-100) with protease inhibitor and 20 μl Dynabeads Protein G for pre-clearing. After preserving 1% as an input sample the pre-cleared chromatin was incubated with 2 μg antibody, 50 μl 10% BSA and 5 μl tRNA (10 mg/ml) overnight at 4 °C with rotation. Immune complexes were retrieved with 20 μl BSA blocked Dynabeads Protein G for 2 h and washed 2x with RIPA-LS, 2x with RIPA-HS (10 mM Tris-HCl, pH 8.0, 500 mM NaCl, 1mM EDTA, pH 8.0, 0.1% SDS, 0.1% Na-Deoxycholate, 1% Triton X-100), 2x with RIPA-LiCl (10 mM Tris-HCl, pH 8.0, 250 mM LiCl, 1mM EDTA, pH 8.0, 0.5% IGEPAL, 0.5% Na-Deoxycholate) and 1x with 10 mM Tris, pH 8.0. Libraries were then prepared directly on the bead-bound chromatin according to a ChIPmentation procedure that is optimized for low cell number samples ^17^. Briefly, beads were resuspended in 25 μl of tagmentation reaction buffer (10 mM Tris, pH 8.0, 5 mM MgCl_2_, 10% v/v dimethylformamide) containing 1 μl Tagment DNA Enzyme from the Nextera DNA Sample Prep Kit (Illumina) and incubated at 37°C for 10 min followed by 2x washing in RIPA-LS and TE (10mM Tris-HCL, pH 8.0, 1mM EDTA, pH 8.0). The DNA was then de-crosslinked and eluted from the beads in ChIP elution buffer (10mM Tris-HCL, pH 8.0, 5mM EDTA, pH 8.0, 300mM NaCl, 0.4% SDS) with proteinase K at 55 °C for 1 h and 65 °C overnight. ChIP DNA was purified with Qiagen MinElute kit (QIAGEN) and enrichment cycles for library amplification was assessed by qPCR. The libraries were PCR amplified with KAPA HiFi Hotstart Ready mix (Sigma-Aldrich) and barcoded Nextera custom primers ^17^ and finally size-selected (250–350 bp) using Agencourt AMPure XP beads (Beckman Coulter) and validated using the Agilent High Sensitivity DNA Kit with Agilent 2100 Bioanalyzer. Equimolar quantities of libraries were combined for multiplexing to obtain 40 million reads per library. ChIP-seq libraries were sequenced using a NextSeq platform with single-end reads of 75 bases.

### ChIP-seq alignment and peak calling

ChIP-seq reads were aligned to the mouse genome (*M. musculus*, UCSC mm10) using Bowtie2 followed by exclusion of mapping quality scores < 30 by Samtools (v1.2) ^18^ and removal of duplicate reads with Picard (v2.6.0) and blacklisted regions with Bedtools (v2.13.3) ^19^. The processed BAM files were converted to bigwig files using Deeptools. Peaks for histone marks were called with MACS2 (v.2.1.0) with settings: -- extsize=300 --q 0.05 --keep-dup all --nomodel --broad and for Utx and transcription factors with MACS1.4 (v1.4.2) using default parameters and *P*<10^−10^. Enriched regions were scored against matching input libraries. All experiments were done as biological replicates. For histone marks and UTX ChIPs only enriched regions that overlap in replicates were retained as consistent peaksets. Publically available datasets were processed identically. For HNF1A ChIPs, peaks from two replicates that overlap by at least one base were merged and replaced with a single peak. Library information of public and internal datasets is provided in **Supplementary Table 8**.

### Integrative analysis

Active promoters were defined as consistent H3K27ac peaks occurring within 1kb of an annotated transcription start site (GENCODE GRCm38.p5) and all remaining peaks were considered as active enhancers. HNF1A and UTX peaks were annotated with HOMER (v4.10.3) ^20^ to define TSS-proximal, TSS-distal, intronic, exonic, and intergenic regions, and were assigned to genes using the “closest” function in Bedtools with default parameters. Fisher’s exact test in R was used to define enrichment of HNF1A- and UTX-bound regions and active promoter and enhancer regions, or differentially expressed genes in the *Hnf1a^aKO^* and *Utx^pKO^* datasets. Expressed genes (Log2 TPM>2.5) were used as background. Aggregation plots were calculated with Deeptools. Briefly, all Utx bound regions were extended ±10 kb or ±5 kb from the center of the peaks for analysis of H3K27me3 or H3K27ac signals, respectively. The resulting regions were then divided into 10 bp bins and the mean value in each bin was calculated based on the normalized values given in the bigwig files for the histone marks. To assess the relationship between functional UTX binding sites and UTX-dependent histone modification changes, the Log_2_ fold change of the averaged normalized H3K27me3 signals were plotted against the Log_2_ fold change of H3K27ac signals in each Utx-bound region. Differential binding of UTX and HNF1A was analyzed with DiffBind (v2.6.6) ^21^ using the DEseq2 method without control input read counts. Differential peaks with FDR<0.05 were defined as significant.

### Functional Annotation and Gene Set Enrichment Analysis (GSEA)

The ENRICHR tool ^22, 23^ was used to functionally annotate differentially expressed genes. Significantly enriched signatures were identified using Bonferroni corrected *P*<0.01. The GSEA pre-ranked analysis ^24^ was used to determine whether predefined sets of genes show significant concordant differences with gene expression changes in RNA-seq datasets. For GSEA we used gene sets from the Molecular Signature Database ^25^, custom made sets of differentially expressed genes from our dataset, gene sets of pancreatic cell types ^26^, and human PDAC ^27–29^. All were analyzed with the parameters: numbers of permutations = 10000 and scoring scheme = weighted.

### Motif Analysis

To find enriched *de novo* transcription factor DNA-binding profiles we used HOMER ^20^ and searched for 6, 8, 10, and 12 bp sized motifs in ±250bp regions surrounding the center of Utx -bound regions within promoters and enhancers that were functionally associated with transcriptional changes. As background sequences we used all H3K27ac regions. DNA-motifs were annotated with HOMER and retrieved if the HOMER score was more than 0.7. Centrimo from the MEME suite ^30^ was used to compute enrichment of known HNF1A motifs in HNF1A-bound regions (**Supplementary Figure 2g**) and in UTX-bound regions (**Figure 7d**). Position weight matrices for HNF1 were retrieved from JASPAR ^31^ (http://jaspar.genereg.net/) and HOCOMOCO ^32^ (http://hocomoco11.autosome.ru/) databases.

### Data Visualization

Bigwig files from RNA-seq and DNA-seq experiments were visualized in the Genome Browser (http://genome.ucsc.edu/). R packages were used to create boxplots (BoxPlotR) ^33^, bar and scatter plots (ggplot2) ^34^. Enrichment plots from GSEA were drawn by Genepattern ^35^. Aggregation plots and heatmaps of ChIP experiments were drawn with Deeptools. GraphPad Prism 6 was used for bar graphs, calculation of contingency tables and piecharts. To draw heatmaps we used Morpheus (https://software.broadinstitute.org/morpheus).

### Analysis of human PDAC genomic data

#### Data acquisition and preprocessing

RSEM normalized RNA-seq data from the TCGA-PAAD cohort was downloaded from the Firehose browser (https://gdac.broadinstitute.org/). Molecular subtypes and purity class was assigned to each sample as defined by the Cancer Genome Atlas Research Network ^36^. Samples with high purity (76 samples) were retained for further analysis. Quantile normalized background adjusted array expression data from the ICGC-PACA-AU cohort was downloaded from the ICGC data portal (https://dcc.icgc.org/). Probe sets (ILLUMINA HumanHT 12 V4) were assigned to gene names. Multiple probe sets mapping to the same gene were collapsed using the collapseRows algorithm from the WGCNA package ^37^ keeping the gene-probe set combination with highest variance across all samples. Purity class was assigned to each sample according to Bailey et al. ^27^ retaining 121 high purity samples for further analysis.

#### Consensus clustering

Samples from the ICGC-PACA-AU study were classified into molecular subtypes based on signature genes from Collisson et al. ^28^, Moffitt et al. ^29^, or Bailey et al. ^27^. Out of the 62 PDA assigner genes identified by Collisson et al., 57 matched with our preprocessed data. We also selected 50 genes with the highest gene weights for the basal and classical PDAC subtypes (100 genes in total) from Moffitt et al’s gene factorization analysis, out of which 94 genes matched with our data. From Bailey et al.’s 613 differentially expressed genes across their defined classes (Squamous, ADEX, Immunogenic, and Pancreatic Progenitor), we retained 457 matching genes. We then applied consensus clustering to the ICGC-PACA-AU data with the signature genes from the three studies using ConsensusClusterPlus (CCP) v.1.24.0 ^38^. Each gene expression profile was first Log_10-_ transformed and median centered. We next performed 1000 iterations of CCP using Pearson correlation as the distance metric, partitioning around medoids, and a random gene and sample fraction of 90% in each iteration. We verified that the obtained groups and their gene expressions reflected the up and down relationships for each gene in each group described in the three studies. We next used human orthologs of differentially expressed genes from our study in *HNF1A*-deficient mouse pancreas to cluster the TCGA-PAAD and ICGC-PACA-AU cohorts. We applied consensus clustering with non-negative matrix factorization, using Pearson distance metrics and 2000 descent iterations ^39^, and thereby identified a group of samples with similar expression signature to the *Hnf1a^aKO^* pancreas. To generate expression heatmaps, we identified genes that were differentially expressed across the clusters (FDR<0.05) with a SAM multiclass analysis (samr v.2.0) ^40^, Z-score transformed each row of the matrix, then used Morpheus (https://software.broadinstitute.org/morpheus) to cluster only the rows, using a one minus Pearson correlation distance metric, average linkage method and Hierarchical clustering.

To identify tumors with most pronounced HNF1A-deficient function, we first defined a gene set containing 106 human orthologs of HNF1A-bound genes that were downregulated in *Hnf1a^aKO^* mouse pancreas. We then interrogated that behavior of this gene set in every tumor sample. This was carried out by GSEA (see also GSEA subheading), using the HNF1A-dependent gene set, and testing the enrichment in each tumor’s gene lists rank-ordered by differential expression of all genes in the tumor vs the median expression of all genes across all samples. This analysis revealed 3 groups of tumor samples: (1) HNF1A loss-of-function (LoF) with NES<0, P<0.05, (2) *Control 1* with NES<0, P>0.05, and (3) *Control 2* with NES>0.

#### UTX mutations

Information on *UTX* mutations was retrieved from the ICGC data portal (https://dcc.icgc.org) and Bailey et al. ^27^. 15 samples in the high-purity ICGC-PACA-AU cohort had *UTX* mutations, which we separated into 3 groups according to mutation types defined by ICGC: (1) Deletions (small <=200bp), (2) Substitutions (single base), (3) Insertions (small <=200bp. The fourth group, (4) Structural Variants, were from Bailey et al. Mutations defined by ICGC as having ‘high’ functional impact in (1-3) and having a ‘loss of function’ consequence in (4) were all frame-shift mutations. We considered those mutations as being functional deleterious and defined samples with such genetic alterations in *UTX* as *UTX* loss-of-function (LoF) tumors. The rest were defined as samples with *UTX* mutations of unknown consequence. Significantly Mutated Genes (SMGs) were from Bailey et al.’s ^27^ SMG analysis using Intogen, Mutsig and HOTNET. Genes considered significantly mutated were significant in >1 analysis.

### Data deposition

RNA-seq and ChIP-seq data sets generated here are available in the ArrayExpress database at EMBL-EBI (www.ebi.ac.uk/arrayexpress) under accession numbers E-MTAB-7944 and E-MTAB-7945.

## Supplementary Figure Legends

**Supplementary Figure 1.**
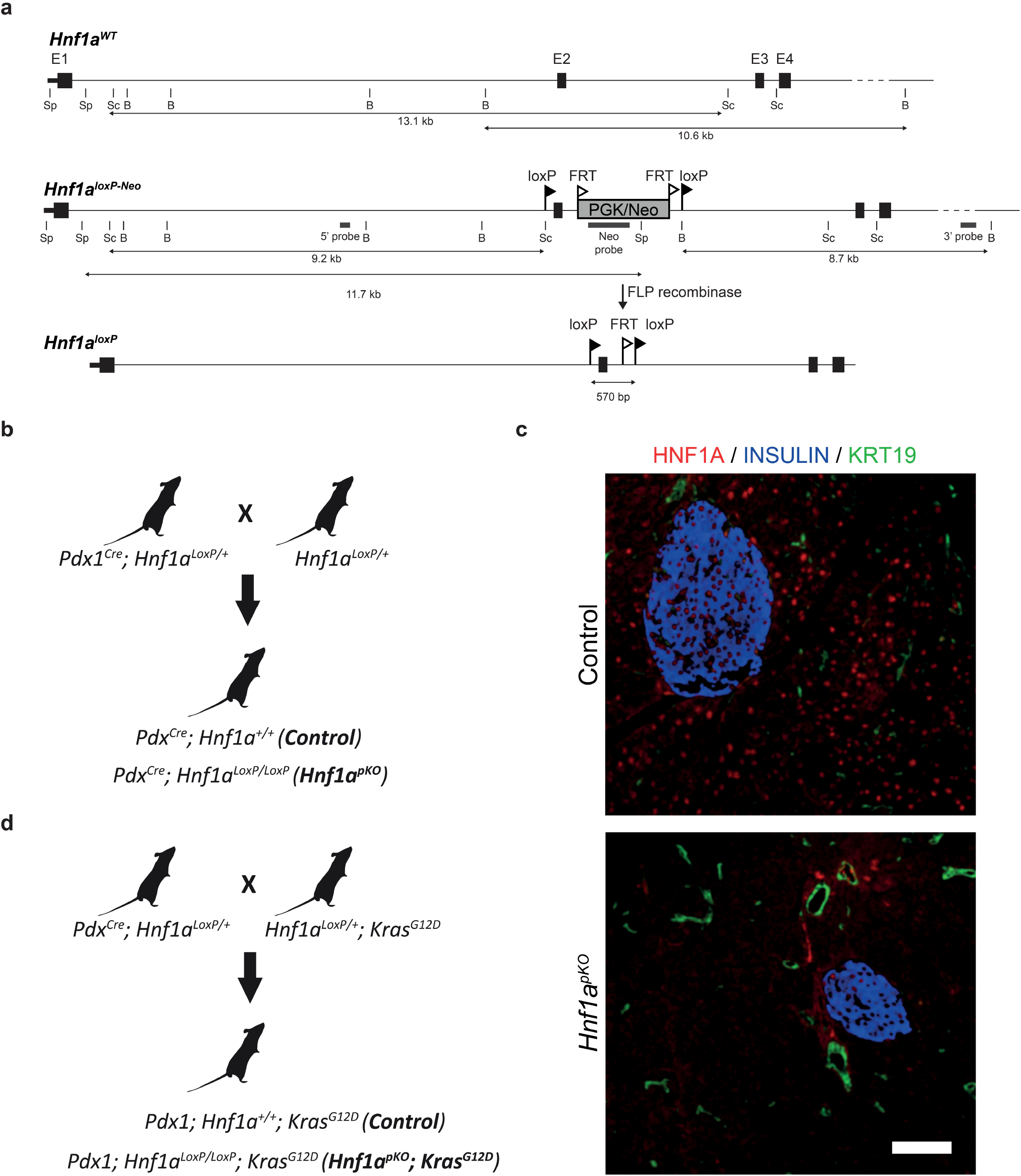
Strategy for creating a conditional Hnf1a mutant mouse line expressing Kras^G12D^.

**a**, Schematic of the targeted *Hnf1a* locus and the resulting *floxed exon 2* alleles, before and after excision of the PGK/Neomycin cassette by FLP-FRT recombination. **b**, Strategy for generating *Hnf1a^pKO^* and control *Pdx1^Cre^*; *Hnf1a*^+/+^ mice. **c**, Immunofluorescence analysis shows that HNF1A (red) was efficiently excised in most acinar and endocrine cells from adult *Hnf1a^pKO^* mice. Scale bar represents 50 μm. d, Strategy for generating *Hnf1a^pKO^*;*Kras^G12D^* and control *Pdx1^Cre^*;*Hnf1a^+/+^*; *Kras^G12D^* mice. B, BamHI; Sc, ScaI; Sp, SpeI.

**Supplementary Figure 2.**
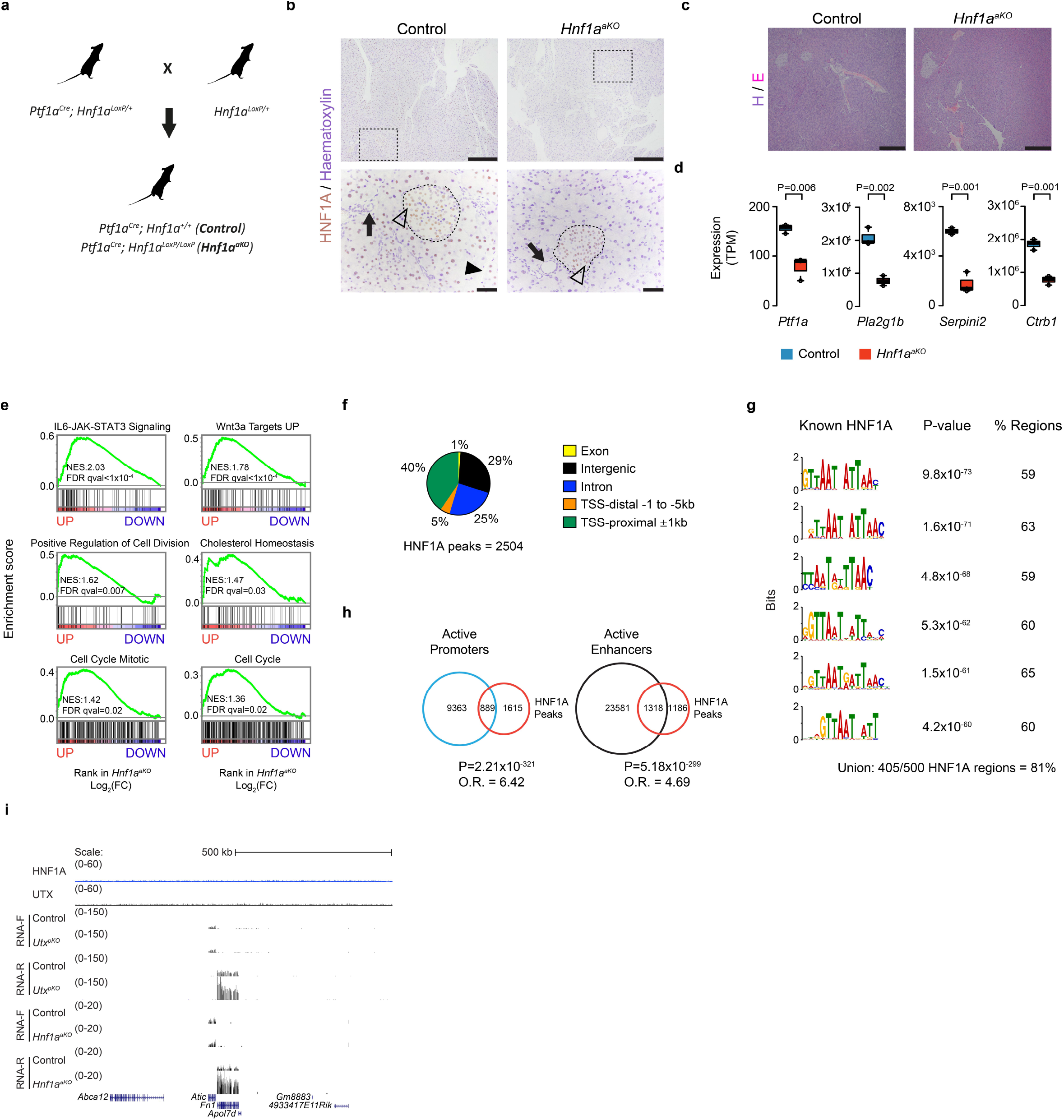
Hnf1a deletion leads to abnormal acinar cell identity and increased expression of genes related to oncogenic pathways. **a**, Breeding strategy to generate *Hnf1a^aKO^* and *Ptf1a^Cre^*;*Hnf1a*^+/+^ control mice using *Ptf1a^Cre^* and *Hnf1a^LoxP^* alleles. **b**, *Ptf1a^Cre^* deletes HNF1A efficiently in acinar cells but to a lesser extent in islets of Langerhans. HNF1A IHC and haematoxylin staining in pancreas of control and *Hnf1a^aKO^* mice. HNF1A is expressed in acinar and islet cells, but not in ductal cells in normal pancreas (left). HNF1A expression is depleted in acinar cells but largely not in islets in *Hnf1a^aKO^* pancreas (right). The squared dotted boxes (top) indicate magnified areas (bottom). Arrows point at ducts, arrow head at HNF1A positive acinar cell, and open arrow head at HNF1A positive islet cell. The dotted encircled areas indicate islets of Langerhans. Scalebar (top) 300μm, (bottom) 50μm. **c**, H&E stainings in pancreas of control (left) and *Hnf1^aKO^* mice (right) showing unaltered pancreatic morphology. Scale bar 300μm. **d**, Expression of acinar differentiation genes in pancreas from *Hnf1a^aKO^* and controls, depicted as box plots with median values and interquartile range of TPM values. P-values were determined by two-tailed Student’s t tests. **e**, GSEA showing increased expression of oncogenic pathways in *Hnf1a^aKO^* pancreas. **f**, Distribution of pancreatic HNF1A binding sites in annotated genomic regions. **g**, Enrichment of known HNF1 motifs in the top 500 most significant HNF1A-bound ChIPseq regions and percentage of regions containing each motif. The “union” is the percentage of regions with at least one motif sequence occurrence. Enrichment P-values are calculated using the one-tailed binomial test. **h**, Venn diagrams illustrating that HNF1A-bound regions are enriched in regions of active promoters and enhancers. P-values and odds ratios were calculated by Fisher’s exact test. **i**, Genome browser track for the *Fibronectin* (*Fn1*) gene showing upregulated expression in *Hnf1a^aKO^* pancreas, and absence of HNF1A or UTX binding to its locus.

**Supplementary Figure 3.**
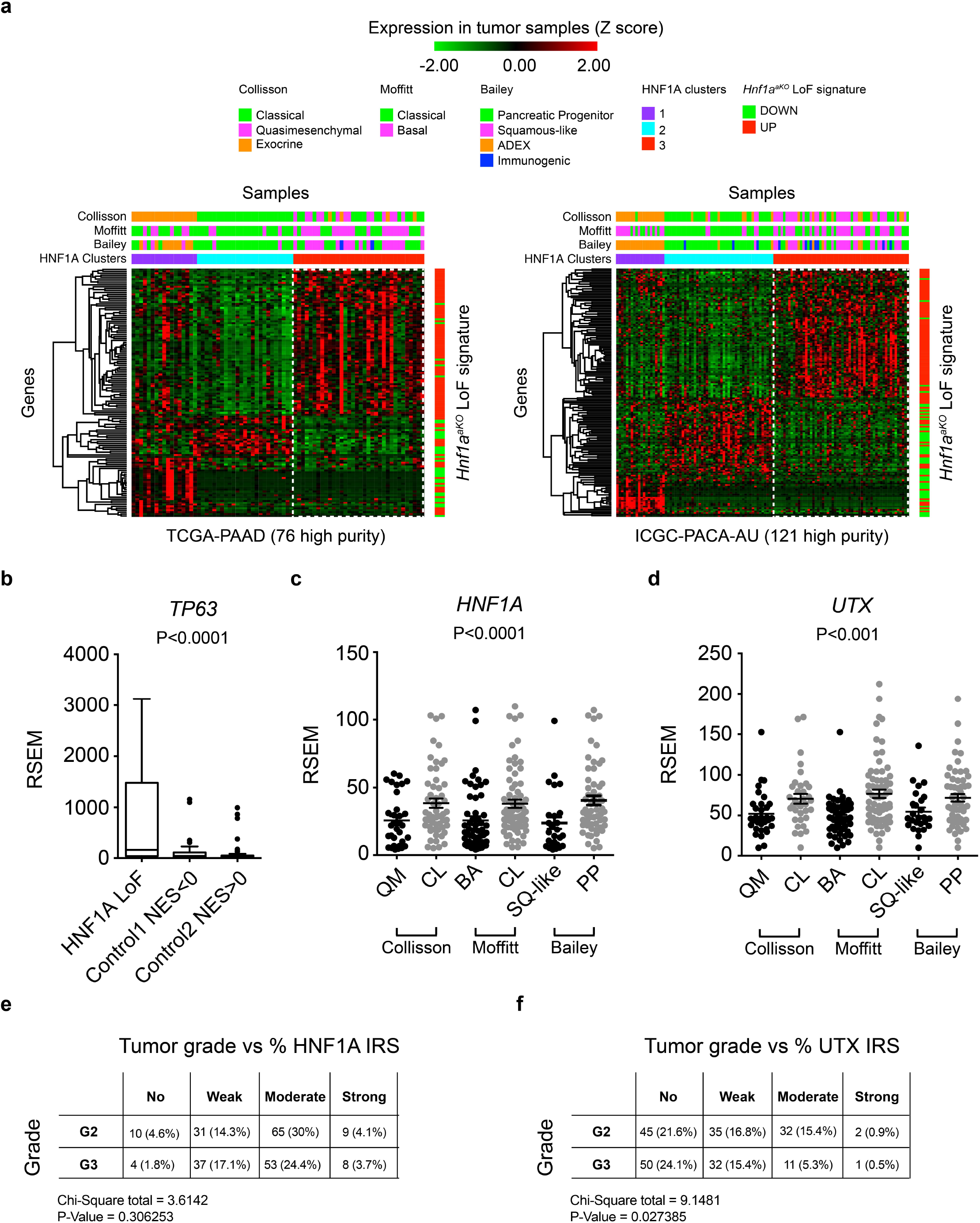
HNF1A signature correlates with non-classical PDAC subtypes. **a**, Consensus clustered Z score-normalized gene expression heatmaps of high-purity TCGA-PAAD and ICGC-PACA-AU human PDAC samples. Clustering was performed with nonnegative matrix factorization based on expression of significantly down- and upregulated genes in *Hnf1a^aKO^* pancreas. This revealed a cluster (HNF1A cluster 3) with concordant up- and downregulation of genes in *Hnf1a^aKO^* pancreas, which predominantly matched non-classical PDAC molecular subtypes (quasimesenchymal, basal, squamous-like, pink in top tracks), as opposed to classical PDAC subtypes (green in top tracks). Multiclass SAM differentially expressed genes (q<0.05) between HNF1A clusters are shown. Genes were hierarchically clustered using complete linkage with one minus Pearson correlation metrics. Along the right side of the heatmaps are green and red indicators of down- and upregulated genes in *Hnf1a^aKO^* pancreas, respectively. **b**, *TP63* expression was increased in HNF1A LoF tumors compared to control tumors. RSEM normalized count data are shown as box plots with interquartile range, median and whiskers. **c-d**, RSEM normalized expression of *HNF1A* and *UTX*, showing downregulation in non-classical PDAC subtypes (P, Kruskal-Wallis). **e,f**, HNF1A levels are not lower in high histological grade PDAC (e) while UTX levels are (f). To determine if histological grade of human PDAC was associated with expression levels of HNF1A (**e**) or UTX (**f**) proteins, we evaluated contingency tables of tumor grades versus staining intensities of each case in tissue microarray (TMA) IHC. Tumor grades were scored as either moderately differentiated (G2) or poorly differentiated/high grade (G3) and staining intensities were expressed as an Immuno Reactivity Score (IRS) reflecting either No, Weak, Moderate, or Strong staining intensities (see material and methods for details). Numbers of cases and percentages (in brackets) out of total cases are indicated for each tumor grade and staining intensity. The Chi-squared test was used to determine the probability of a significant relationship. Chi-square and P-values are shown. N=217 patients for HNF1A and N=208 patients for UTX.

**Supplementary Figure 4.**
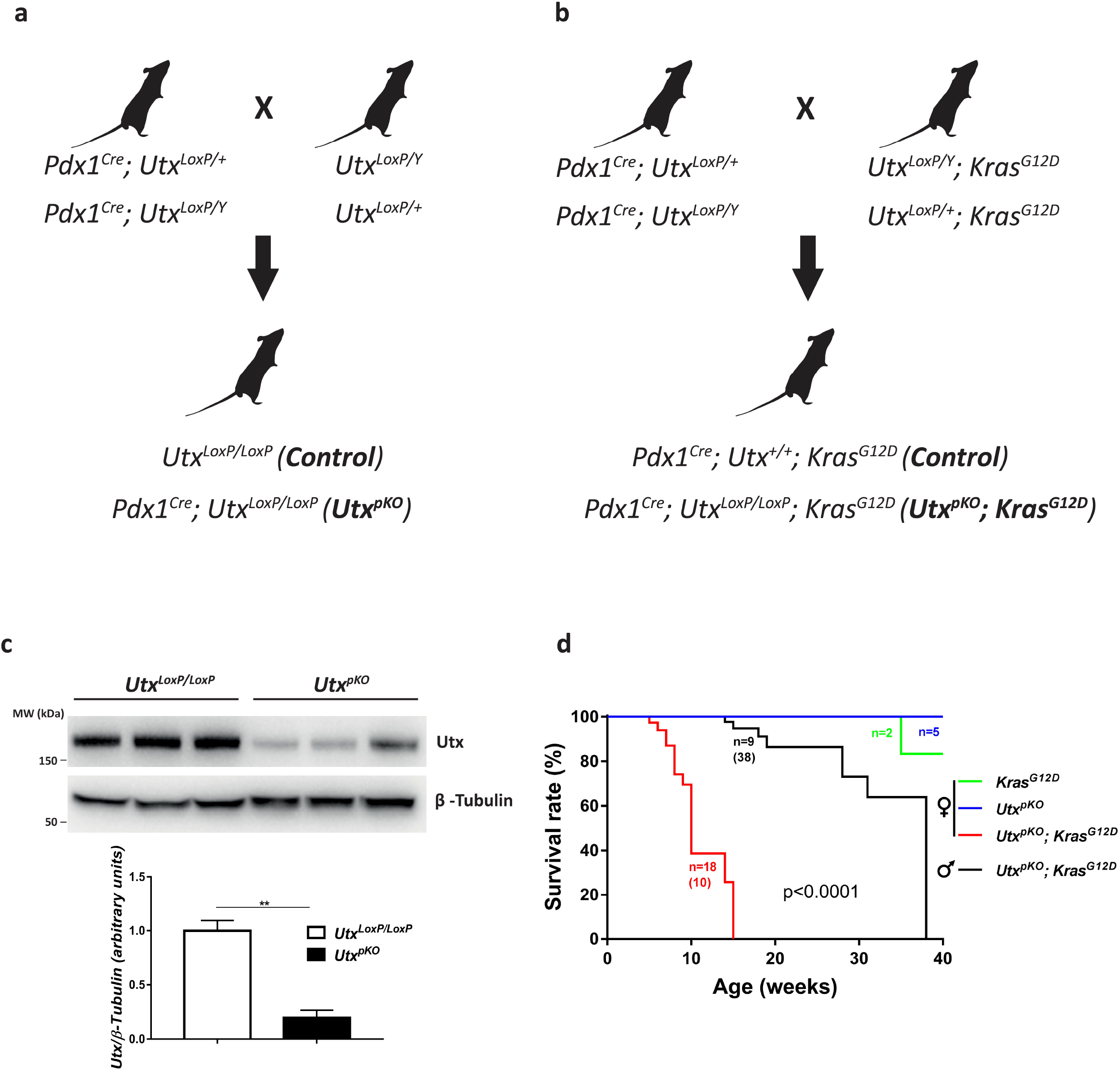
Pancreatic Utx mutant mice expressing Kras^G12D^. **a,b**, Schematic of strategy for generating *Utx^pKO^*;*Utx^pKO^*;*Kras^G12D^* and control mice. **c**, Immunoblot showing reduced expression of UTX and average quantification of UTX relative to β-tubulin in pancreas from *Utx^LoxP/LoxP^* and *Utx^pKO^* mice, ** p<0.01, Student’s *t* test. **d**, Kaplan-Meier plot showing the survival of female *Kras^G12D^, Utx^pKO^* and male and female *Utx^pKO^*;*Kras^G12D^* mice, n= number of mice and numbers in brackets show median survival. Group median survival shown in brackets was compared using the Log-rank (Mantel-Cox) test.

**Supplementary Figure 5.**
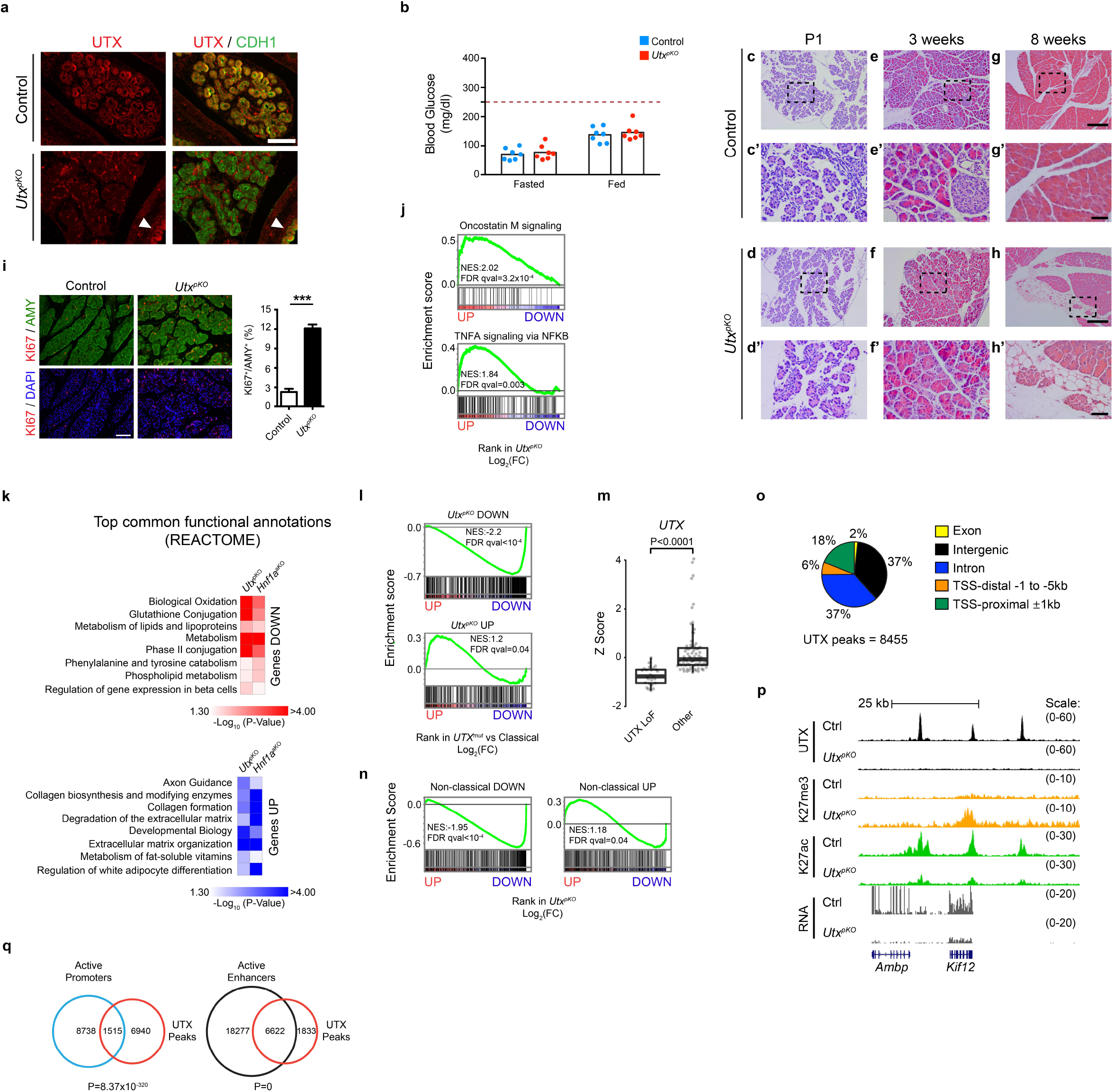
UTX Suppresses Growth and Oncogenic Pathways and Maintains Acinar Cell Integrity. **a**, Efficient deletion of *Utx* in the pancreatic epithelium at E15.5. UTX (red) is ubiquitously expressed in all pancreatic cells. CDH1 (green) marks epithelial cells. Upon deletion, UTX staining is lost specifically in CDH1-expressing epithelial cells but not in mesenchymal cells or in the stomach epithelium (white arrow heads). **b**, *Utx* mutant mice show normal fasting and fed glycemia. The horizontal stroked line indicates blood glucose levels at 250mg/dl as a reference for levels indicative of diabetes mellitus. **c-h**, The pancreas of *Utx^pKO^* mice were histologically normal until 8 weeks of age, at which point there was occasional acinar cell attrition. Scalebars, 250 μm (10x magnification), 50 μm (40x magnification). **i**, Representative picture (left) showing increased number of KI67 (red) amylase-expressing acinar cells (green) in *Utx^pKO^* pancreas. Scalebar, 250 μm. Quantifications (right) were performed on three pancreatic sections separated by at least 100μm from 4 control and 4 *Utx^pKO^* mice. **j**, GSEA plots showing enrichment of Oncostatin M and “TNFA signalling via NFKB” gene sets among genes upregulated in *Utx^pKO^* pancreas. **k**, Examples of most significantly deranged REACTOME pathways in both UTX and HNF1A-deficient pancreas (see also **Supplementary Table 5**). **l**, *Utx^pKO^* down and upregulated gene sets showed concordant deregulation in *UTX* LoF mutant tumors vs classical PDAC based on Bailey et al’s signature. Normalized enrichment scores (NES) and FDR q-values are shown. **m**, Tumors with UTX-deficient phenotypes showed decreased *UTX* mRNA. We created a gene set of human orthologs of *Utx^pKO^* downregulated genes, and for each high purity tumor sample in the ICGC-PACA-AU, we used GSEA to test for enrichment of this gene set in gene lists that were rank-ordered by differential expression in the individual sample vs. all other samples. Samples with NES<0 and P-value<0.05 were considered as having UTX LoF phenotypes and were compared against all other samples. Z-score normalized count data are shown as box plots with interquartile range, median and whiskers. P-values were determined by two-tailed Student’s t test. **n**, Gene sets that showed up- or downregulation in non-classical human PDAC showed concordant enrichment in up- or downregulated genes in *Utx^pKO^* vs control pancreas. GSEA NES and FDR q-values are shown. **o**, Genomic distribution of UTX binding sites in mouse pancreas. **p**, Genome browser track of UTX, H3K27me3 and H3K27ac ChIPseq, and RNAseq in the genes *Ambp* and *Kif12* in control and UTX-deficient pancreas. **q**, UTX-bound regions are enriched in active pancreas promoters and enhancers. P-values are calculated by Fisher’s exact test.

**Supplementary Figure 6.**
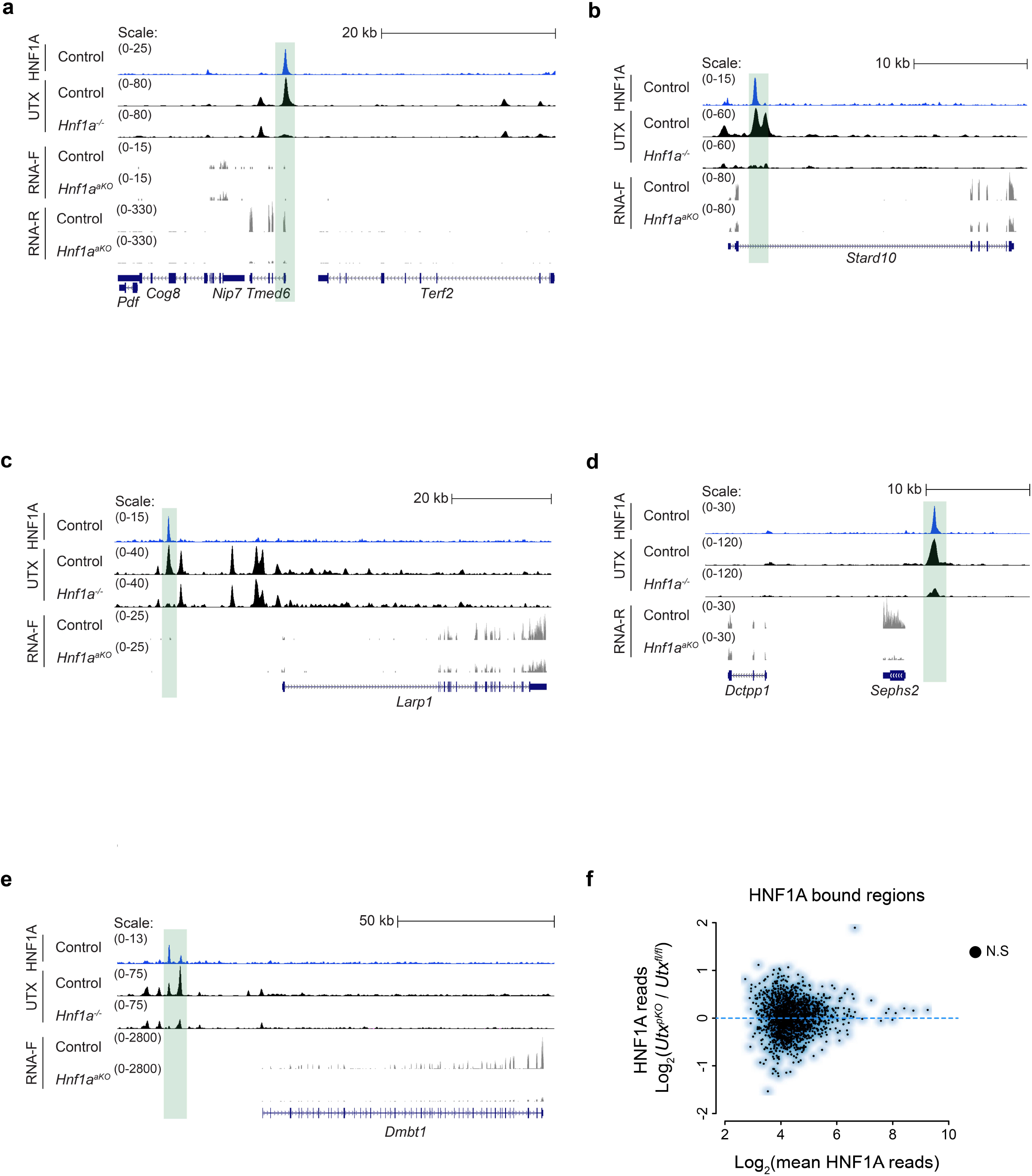
UTX binding to target genes is dependent on HNF1A while HNF1A can bind to chromatin in the absence of UTX. **a-e**, Examples of loci co-bound by HNF1A and UTX, showing loss of UTX binding in HNF1A-deficient pancreas (highlighted in green), and decreased RNA levels in HNF1A-deficient pancreas. **f**, HNF1A binding to chromatin is unaffected in *Utx^pKO^* pancreas.

## Supplementary Tables

**Supplementary Table 1.** Differential expression *Hnf1a^aKO^* pancreas

**Supplementary Table 2.** Functional annotation of *Hnf1a^aKO^* pancreas

**Supplementary Table 3.** Differential expression in *Utx^pKO^* pancreas

**Supplementary Table 4.** Functional annotation of *Utx^pKO^* pancreas

**Supplementary Table 5.** Shared annotations of *Hnf1a^aKO^* and *Utx^pKO^* pancreas

**Supplementary Table 6.** Genotyping primers

**Supplementary Table 7.** Antibodies list

**Supplementary Table 8.** RNA-seq and ChIP-seq datasets

